# Mice with mono-allelic p.R37H *Dhdds* variant show aberrant glycosylation and interneuron deficits

**DOI:** 10.1101/2025.08.15.670547

**Authors:** Afitz Da Silva, Merrick S. Fallah, Samuel Boris Tene Tadoum, Mehrnaz Fazeli, Irena J.J. Muffels, Siyan Wang, Francois Grenier, Rohit Budhraja, Luisa Sturiale, Angela Messina, Moshe Giladi, Mahsa Taherzadeh, Éric Bonneil, Shaukat Khan, Danielle te Vruchte, Yojiro Yamanaka, Graziella Di Cristo, Fadi F. Hamdan, Frances M. Platt, Shunji Tomatsu, Yoni Haitin, Tamas Kozicz, Pierre Thibault, Domenico Garozzo, Akilesh Pandey, Eva Morava, Elsa Rossignol, Alexey V. Pshezhetsky

**Author notes:** Correspondence to: Alexey V. Pshezhetsky, CHU Sainte-Justine, 3175 Côte Ste-Catherine, Montréal (QC) H3T 1C5 – Canada, Elsa Rossignol, CHU Sainte-Justine, 3175 Côte Ste-Catherine, Montréal (QC) H3T 1C5 – Canada. These authors contributed equally to this work (Merrick S. Fallah, Samuel Boris Tene Tadoum, Mehrnaz Fazeli). Equally contributed as senior authors (Elsa Rossignol, Alexey V. Pshezhetsky).

## Abstract

Developmental delay and seizures with or without movement abnormalities (OMIM 617836) caused by heterozygous pathogenic variants in the *DHDDS* gene (DHDDS-CDG) is a rare genetic disease that belongs to the progressive encephalopathy spectrum. It results in cognitive delay in affected children, accompanied by myoclonus, seizures, ataxia and tremor, which worsens over time. *DHDDS* encodes a subunit of a DHDDS/NUS1 cis-prenyltransferase (*cis-*PTase), a branch point enzyme of the mevalonate pathway essential for N-linked glycosylation. We describe the first mouse model of this disease, *Dhdds^R37H+/-^*strain, heterozygous for the human recurrent *de novo* c.110G>A:p.R37H pathogenic variant. *Dhdds^R37H+/-^* mice present with seizures, myoclonus and memory deficits associated with reduced density or/and maturity of inhibitory interneurons in the cortex. Multiomics analyses of mouse CNS tissues, together with the enzymatic/structural characterization of the R37H DHDDS mutant protein, reveal that the variant produces a catalytically inactive enzyme and results in a brain dolichol deficit, aberrant glycosylation of brain glycoproteins, including those involved in synaptic transmission and major perturbations in the CNS proteome and lipidome. Acetazolamide, a carbonic anhydrase inhibitor clinically approved for treatment of glaucoma, epilepsy, and intracranial hypertension, and successfully used “off-label” to treat genetic movement disorders, reduces seizure susceptibility to pentylenetetrazol in *Dhdds^R37H+/-^* mice, suggesting potential therapeutic value of using this drug in human DHDDS-CDG patients. Together, our results define *cis-*PTase as a master regulator of CNS development and function and establish that its monoallelic debilitating variants cause a novel congenital disorder of glycosylation associated with aberrant levels of neuronal proteins and lipids.

## Introduction

Protein glycosylation plays a crucial role in the development and function of the central nervous system, regulating essential processes including axon pathfinding, neurite outgrowth, synaptogenesis, neurotransmission, and many others.^1^ It is not surprising, therefore, that disruptions or alterations in this process in rare genetic diseases known as Congenital Disorders of Glycosylation (CDG) lead to severe physiological consequences with over 80% of patients presenting with neurologic symptoms including seizures, movement disorders and intellectual disability.^2^ Identification of novel CDG has exploded over the past decade with ∼200 types characterized to date. CDG, collectively, affects most glycosylation pathways, including N- and O-glycosylation of proteins, synthesis of glycosphingolipids, and formation of GPI-anchors.^3^ In many cases, despite the known genetic defects, the molecular mechanisms underlying clinical manifestations in neurologic CDG are not well understood, and patients exhibiting these symptoms are rarely responsive to currently available treatments.^4,5^ Consequently, no specific therapies exist for the vast majority of CDG subtypes.

Recently, *de novo* heterozygous recurrent Dehydrodolichyl Diphosphate Synthase gene (*DHDDS*) variants have been identified in patients with developmental delay and seizures with or without movement abnormalities. Specifically, the c.110G>A:p.R37H (NM_205861) variant was associated with progressive myoclonus with seizures presenting during the first years of life.^6,7^ DHDDS, together with Nuclear Undercaprenyl Pyrophosphate Synthase 1 (NUS1) forms *cis-*prenyltransferase (*cis-*PTase), the rate-limiting enzyme for dolichol-pyrophosphate biosynthesis, a key metabolic step in the protein N-glycosylation pathway. Both DHDDS and NUS1 *cis-*PTase subunits are required to produce dehydrodolichyl diphosphate, a precursor of dolichol phosphate and its subsequent metabolites, glycosyl carrier lipids, the key molecules for protein glycosylation in the lumen of the endoplasmic reticulum.^8,9^ Previously, a homozygous pathogenic variant p.K42E, adjacent to the catalytic domain of DHDDS, was found in Ashkenazi Jewish patients with retinitis pigmentosa.^10^ The variant is responsible for 8% of retinitis pigmentosa cases in Ashkenazi Jews and 0.8% of cases in the patients with mixed ethnicity. Consecutively, autosomal recessive loss of function *DHDDS* variants were described underlying a lethal form of CDG, DHDDS-CDG.^11^ Recently, however, *de novo* recurrent DHDDS variants were identified in multiple patients with progressive myoclonus, developmental delay and seizures, with or without movement abnormalities. We have reported that the *de novo* c. 110G>A:p.R37H and c.632G>A:p.R211Q variants were associated with intellectual disability and seizures appearing during the first years of life.^6^ Further studies evaluating 25 patients with *de novo* pathogenic variants in *DHDDS*, provided the first systematic description of their clinical features and long-term outcome, and demonstrated that most variants show abolished or severely reduced *cis-*PTase activity, as demonstrated in yeast complementation system and using enzymatic assays.^12,13^ More recently, pathogenic variants in the DHDDS/NUS1 complex have also been detected in over 50 patients with Progressive Myoclonic Epilepsy^14^, demonstrating that the diseases of the DHDDS/NUS1 spectrum are more frequent than initially thought. Surprisingly, in reported patients, in contrast to other CDG involving alterations in dolichol metabolism, hypoglycosylation of serum glycoproteins was not detected and the urinary dolichol levels were normal.^14^ Accumulation of lipidic material and altered lysosomes were detected in the patients’ cultured fibroblasts, suggesting dysfunction of the lysosomal enzymatic scavenger machinery,^6,12,14^ but it was not clear whether these findings extended to the CNS, the organ primarily affected by the disease. Although yeast complementation assays were instrumental in determining the physiological activity of several DHDDS mutants, understanding the disease pathophysiology has been, so far, limited by the predominantly neuronal presentation and the absence of suitable animal models recapitulating human clinical phenotypes.

Here, we describe the first mammalian model of the disease, a knock-in mouse strain (*Dhdds^R37H+/-^*) monoallelic for the recurrent *de novo* R37H human DHDDS variant. This model successfully mimics the key aspects of the human clinical phenotype, including developmental delay and spontaneous seizures, thereby offering a valuable tool for future therapeutic investigations. In *Dhdds^R37H+/-^* mice, this variant leads to a reduction in brain dolichol content associated with abnormal glycosylation and aberrant levels of brain proteins, specifically affecting those critical for the formation and maintenance of synaptic structures. This, together with substantial alterations of brain lipid content, causes neuronal dysfunction and scarcity of interneurons, most likely responsible for the cognitive decline, seizures and myoclonic jerks observed in the mutant mice and human patients.

## Materials and methods

Animal experiments were approved by the Animal Care and Use Committee of CHU Ste-Justine (approval numbers: 2021-3194 and 2021-7105). The knock-in *Dhdds^R37H^* C57BL/6 mouse strain was generated at McGill Integrated Core for Animal Modeling using CRISPR-Cas9 technology^15^ as shown in Supplementary Fig. 1. *Cis*-PTase structural modeling and the molecular dynamics simulations were performed as described, ^16,17^ *cis-*PTase enzymatic activity of the recombinant enzyme and mouse brains microsomes was assayed as described^13,18^ was measured using a radioligand-based assay^13,18,19^. The presence of epileptiform activity in mice was assessed by intracranial video-EEG recordings as described.^20,21^ Seizure susceptibility to pentylenetetrazol (PTZ) was evaluated with a continuous intravenous infusion of the drug as described.^21^ The seizure threshold was defined as the minimal PTZ dose (mg/kg BW) required for seizure induction. Pretreatment with acetazolamide (AZM, 40 mg/kg BW) was conducted 45 min before PTZ infusion.^22,23^ Electrophysiological activity was assessed *in vitro* using voltage-clamp recordings in layer V principal cells from the mouse primary somatosensory cortex. Lipidomic analysis in mouse brain tissues was conducted by LC-MS/MS on UPLC-MRT system (Waters) using CSH C18 Acquity Premier column. Quantification of metabolites was done using Progenesis QI software by integration of areas under chromatograms. MALDI imaging was performed on a MALDI-MRT system (Waters). Mass spectra were integrated using Mass Lynx in negative ion mode at 35 μm pixel resolution. GSLs were analyzed essentially as described.^24^ Analysis of N-linked glycans was performed as previously described^25,26^ by MALDI-TOF and MALDI-TOF/TOF on a 4800 Proteomic analyzer (AB Sciex) in positive polarity and reflector mode.^26^ LC-MS/MS analysis for proteomics and glycoproteomics was carried out on an Orbitrap Eclipse mass spectrometer with Ultimate 3000 liquid chromatography system (Thermo) as described previously.^27,28^ Brain synaptosomes were isolated using a commercially available kit (Thermo #87793) and their protein content analysed as described.^29^ The spontaneous alternation behavior, spatial working memory, exploratory activity, and motor coordination in mice were evaluated using Y-maze, novel object recognition and accelerated rotarod tests, performed as previously described.^15,30,31^ Total RNA sequencing was performed at the Genomics Platform of the Institute for Research in Immunology and Cancer, Montreal. The data are available on: https://amp.pharm.mssm.edu/biojupies/notebook/y4cudhb0o. Analysis of glycans in the bone and brain tissues was conducted as previously described.^32^ Immunofluorescence and lectin fluorescence microscopy of brain cryosections were conducted essentially as described.^33^ The slides were analyzed using a Leica DM 5500 Q confocal microscope and images processed and quantified using ImageJ 1.50i software (National Institutes of Health) in a blinded fashion. For complete methods, see *Supplemental Materials and Methods*.

**Figure 1.**
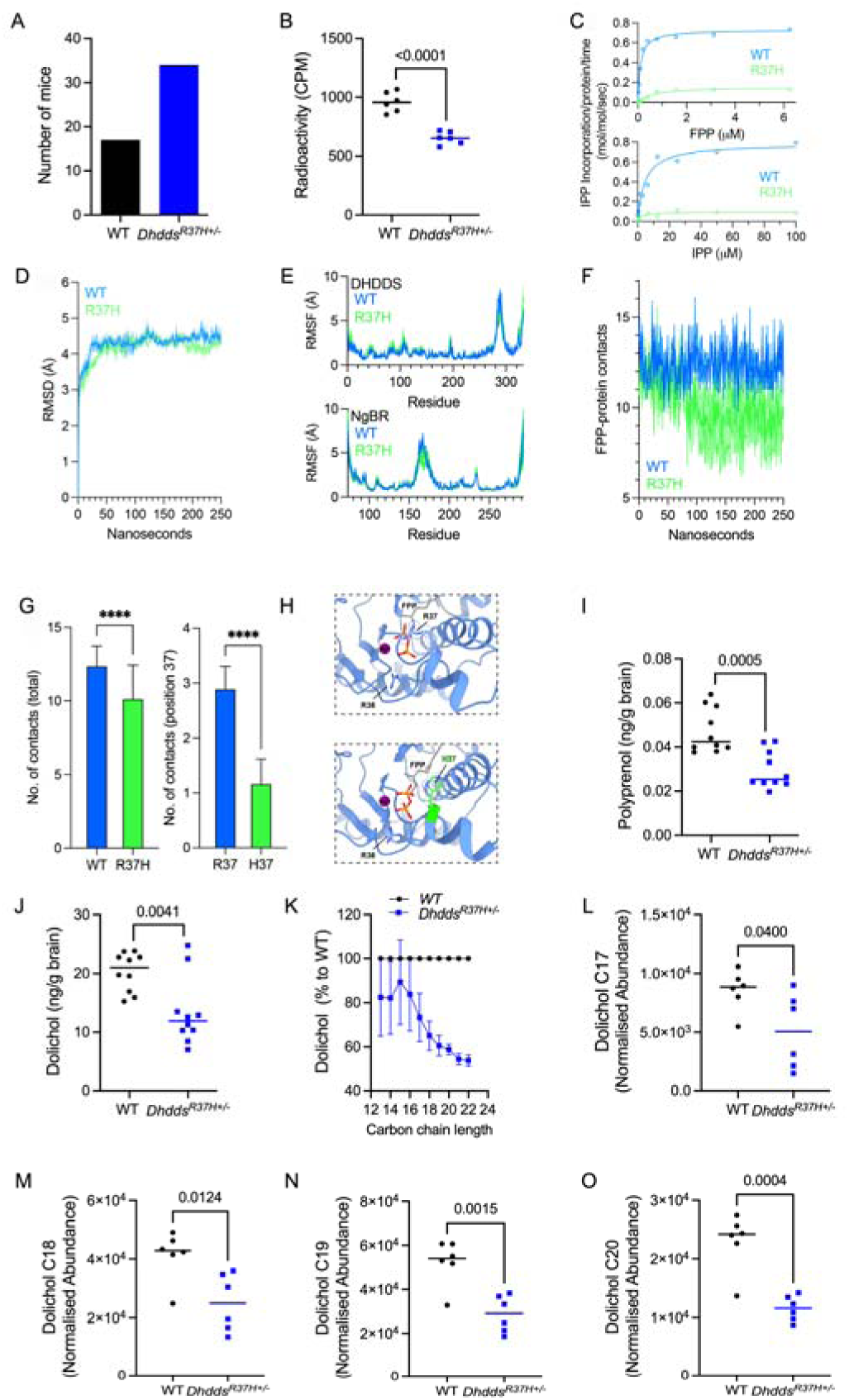
Mouse DHDDS R37H variant impairs *cis-*PTase activity and dolichol production. (**A**) From the offspring of *Dhdds^R37H+/-^* breeding ∼65% of mice are heterozygous for the mutant allele and ∼35%, WT. The results show genotyping of 51 pups. (**B**) *Cis*-PTase activity is reduced in the brain microsomal fraction of *Dhdds^R37H+/-^* compared to WT mice (*n* = 6, 3 males and 3 females for each genotype). (**C**) FPP (upper panel) or IPP (lower panel) dependent purified *cis*-PTase activity was measured as described in the methods section. (**D**) Average root mean square deviation (RMSD) and (**E**) per residue root mean square fluctuation (RMSF) analysis of DHDDS (upper panel) and NgBR (lower panel) for the WT complex (blue) and the complex harboring DHDDS^R37H^ (green) along the simulation trajectory (*n* = 3 for each protein). Shaded area represents the SD. (**F**) Total number of protein-FPP contacts along the simulation trajectories for the WT complex (blue) and the complex harboring DHDDS^R37H^ (green) along the simulation trajectory (*n* = 3 for each protein). Shaded area represents the SD. (**G**) Time-averaged number of protein-FPP contacts for total (left panel) or position 37 (right panel). ****, *P* < 0.0001. (**H**) Representative zoomed-in perspectives on the FPP binding site were obtained from the final simulation frames of the WT (upper panel) or R37H (lower panel). Note the differences in the relative orientations of R37 and H37. (**I-J**) Targeted MS/MS analysis reveals that polyprenols (**J**) and dolichols (**I**) are reduced about two-fold in the brain tissues of *Dhdds^R37H+/-^*compared to WT mice. (**K**) Reduction is more pronounced for the dolichol species with the higher length of the carbon chain (*n* = 10, 5 males and 5 females for each genotype). (**L-O**) Untargeted lipidomic analysis confirms drastic reduction of dolichol C17-C20 species in the brain tissues of *Dhdds^R37H+/-^* mice (*n* = 5, 3 males and 2 females for each genotype). All graphs show individual results, means and SD. *P*-values are calculated by two-tailed *t*-test.

## Results

### Mouse DHDDS^R37H^ variant impairs *cis-*PTase activity and results in reduced dolichol production

The *Dhdds^R37H^* mouse strain was generated as described^34^ and genotyped as shown in Supplementary Fig. 1. In the offspring of *Dhdds^R37H^* heterozygous crosses, ∼65% of mice were heterozygous for the mutant allele and ∼35% were wild-type (WT), consistent with embryonic lethality of homozygous *Dhdds^R37H^*mice (Fig.1A). This was further confirmed by genotyping of >20 E8 embryos in 5 litters from *Dhdds^R37H^* heterozygous breeding, which did not reveal embryos homozygous for the R37H variant.

Enzymatic *cis-*PTase activity, measured in the brain microsomes of heterozygous *Dhdds^R37H+/-^* mice as described,^8,14^ was reduced to 30-40% compared to WT siblings, suggesting that the variant caused a loss of enzymatic activity (Fig. 1B). A previous structural study showed that the R37 residue is highly conserved among *cis*-prenyltransferases and is an essential catalytic residue that directly coordinates the diphosphate group of the farnesyl diphosphate (FPP) substrate within the active site.^13^ Moreover, DHDDS^R37H^, similarly to other variants, DHDDS^G35E^, DHDDS^R37C^, DHDDS^R205Q^ and DHDDS^P233R^, did not support yeast growth and was not able to incorporate radioactive isopentenyl diphosphate (IPP) substrate, suggesting that it is catalytically inactive.^12^ To confirm this, we expressed and purified the recombinant WT and mutated human *cis*-PTase complexes and examined their catalytic activity *in vitro* (Fig. 1C). The complex harboring the DHDDS^R37H^ variant exhibited markedly reduced catalytic activity with *k*_cat_ of 0.15 ± 0.01 s^-1^ compared to 0.73 ± 0.01 s^-1^ for the WT enzyme (*n* = 3 for each protein). Moreover, the R37H variant also increased K_m_ for farnesyl diphosphate, FPP (0.63 ± 0.1 mM vs. 0.1 ± 0.01 mM for the WT enzyme), while not changing the K_m_ for IPP (2.9 ± 3.4 mM vs. 4.5 ± 0.6 mM for the mutant and WT, respectively), consistent with the role of R37H in FPP coordination.

To provide structural insights into the contribution of R37H to substrate coordination, we performed all-atom MD simulations of the FPP-bound complexes (Fig. 1D-H). Each protein was simulated for 250 nsec in triplicate. All the performed simulations have reached convergence, as reflected by the plateau in root mean square deviation (RMSD) values (Fig. 1D).^35^ The per-residue root mean square fluctuation (RMSF) values, representing the spatial fluctuation relative to their average position along the simulation, were similar for the WT or mutant complexes (Fig. 1E). However, inspection of the total number of contacts between the protein and FPP along the simulation trajectories reveals that the complex harboring DHDDS^R37H^ exhibits significantly fewer contacts (Fig. 1F, G). The reduction in the number of contacts (Fig. 1G, left panel) is attributed to the reduced ability of histidine at position 37 to coordinate FPP (Fig. 1G, right panel). This results from the shorter and less positively charged state of the histidine side chain compared to the native arginine (Fig. 1H).

Levels of the *cis-*PTase downstream metabolites, polyprenols and dolichols in the brains of *Dhdds^R37H+/-^* mice, analyzed by targeted liquid chromatography-tandem mass spectroscopy (LC-MS/MS) were markedly (a ∼40% overall reduction compared to WT mice) decreased, consistent with impaired enzymatic activity of the mutant enzyme (Fig. 1I-J). Notably, we also observed a change in the composition of dolichol species, with the most drastic decrease observed for the dolichols bearing the longest (C19-C22) carbon chains (Fig. 1K). These results were confirmed by untargeted lipidomic analysis which revealed a significant and major reduction of C17-C20 dolichol species in the brains of *Dhdds^R37H+/-^* mice (Fig. 1L-O). We did not detect, however, an increase in the levels of the enzyme substrates, farnesyl pyrophosphate (Fig. S2A) and isopentenyl pyrophosphate (Fig. S2B).

As the DHDDS/NUS1 complex is also a branching enzyme of the mevalonate pathway, we analyzed the brain levels of mevalonate/cholesterol synthesis metabolites. Concentrations of squalene and 2,3 oxidosqualene were normal (Supplementary Fig. 2C,D). The mevalonate and mevalonate 5-pyrophosphate levels were unchanged, while mevalonate 5-phosphate was slightly decreased (Supplementary Fig. 2E-G). Acetyl-CoA was slightly increased, but acetoacetyl-CoA and 3-hydroxy-3-methylglutaryl (HMG)-CoA (Supplementary Fig. 2H-J) were also unchanged. Concentrations of cholesterol and other sterol species were similar for *Dhdds^R37H+/-^* and WT siblings with an exception of 7-dehydrocholesterol and dihydrolathosterol that showed an increase in *Dhdds^R37H+/-^*mice (Supplementary Fig. 2K-M).

### *Dhdds^R37H+/-^* mice display memory and neuromuscular deficits, seizures and interictal epileptic activity

In human patients, debilitating heterozygous *DHDDS* variants are associated with a complex, slowly progressive, neurological disorder with a broad spectrum of clinical symptoms including developmental delay, cognitive impairment, abnormal motor control (myoclonus, cortical tremor, ataxia, movement disorder) and epilepsy.^12^ To assess whether these symptoms were recapitulated in the mouse model, we conducted continuous video-EEG recordings for 14 days in *Dhdds^R37H+/-^*and 10 days in WT littermates starting at P60-P75. Electrodes were implanted in the somatosensory cortex (S1) and CA1 of the hippocampus. All *Dhdds^R37H+/-^*mice displayed infrequent (∼1 episode/h) synchronous high amplitude interictal spikes (Fig. 2A-C). Furthermore, spontaneous seizures were detected in 60% (3 out of 5) *Dhdds^R37H+/-^* mice, with a mean seizure frequency of 0.38 episodes/day (*n* = 3) (Fig. 2A-D). The seizures were overall brief and infrequent, with no seizure-related mortality during our recording periods. This was consistent with normal survival of *Dhdds^R37H+/-^* mice. Given the presence of interictal epileptic activity in all mutants, including those that did not present spontaneous seizures, we evaluated the susceptibility to seizure induction with PTZ (10 mg/ml of saline, 0.1 ml/min infusion in the caudal vein) in a separate set of animals. We observed a lower seizure threshold in *Dhdds^R37H+/-^* mice (20.1±0.1 mg/kg BW; *n* = 3) compared to WT littermates (35.2±3.5 mg/kg BW; *n* = 4; *P* = 0.005) (Fig. 2E), confirming an increased seizure susceptibility in mutants.

**Figure 2.**
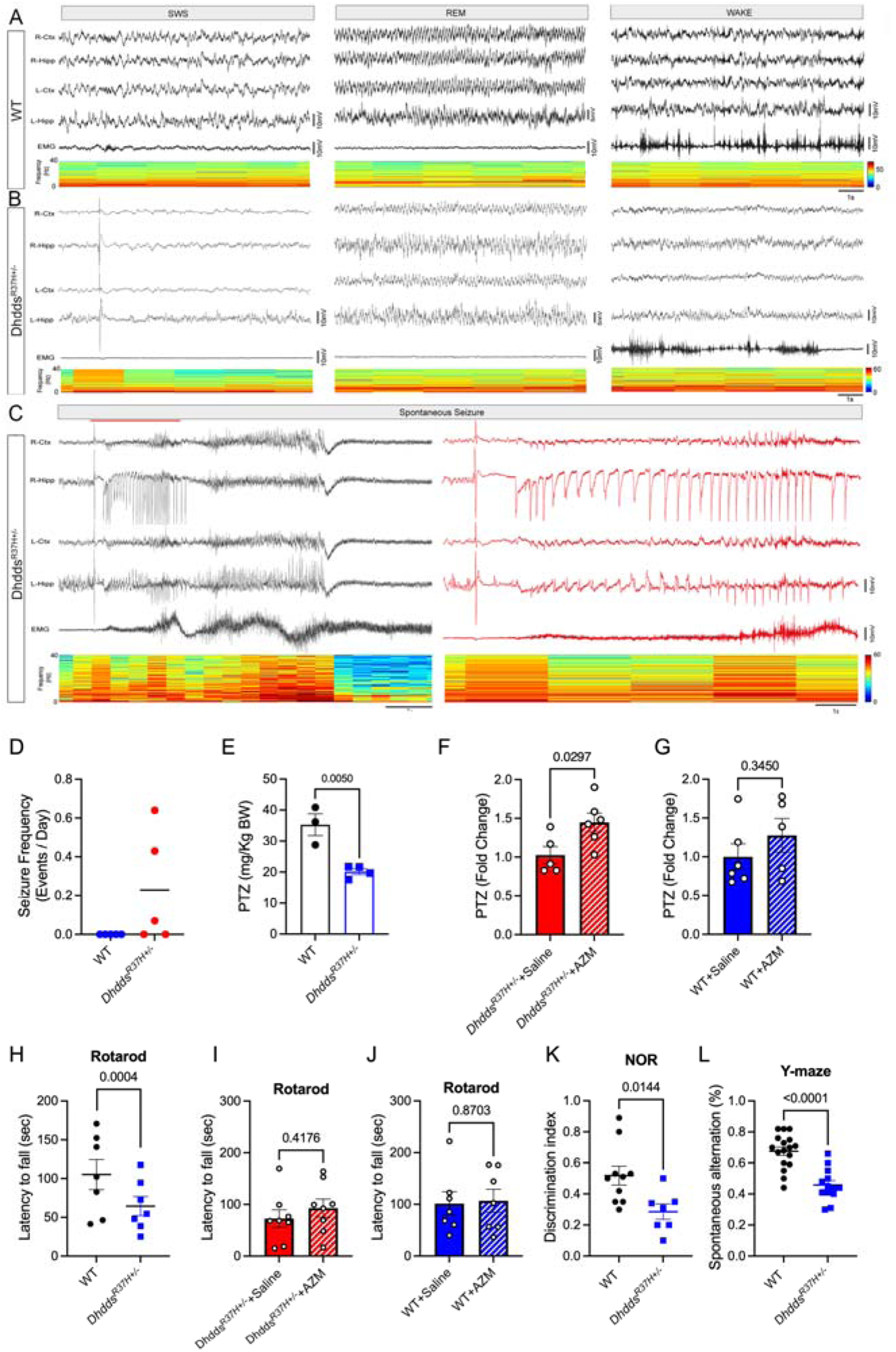
*Dhdds^R37H+/-^*mice recapitulate major aspects of patient’s phenotype. (**A-C**) Illustrative traces of baseline EEG recordings in (**A**) WT controls and (**B**) *Dhdds^R37H+/-^* mice, and (**C**) of a spontaneous seizure recorded in a *Dhdds^R37H+/-^* mouse. Recordings were conducted bilaterally in the somatosensory cortex (S1) and hippocampus (CA1). EEG recordings from *Dhdds^R37H+/-^* mice revealed infrequent interictal spikes. (**D**) Seizure frequency (events/day) in WT and *Dhdds^R37H+/-^*mice. (**E**) Reduced PTZ seizure threshold (minimal dose required for seizure induction) is observed in *Dhdds^R37H+/-^* compared to WT mice (n = 3 WT and 4 *Dhdds^R37H+/-^* mice; *P* = 0.005 Welch’s *t* test). (**F-G**) Single acute injection of acetazolamide (AZM, 40 mg/kg, i.p.), 45-min before PTZ infusion, increases seizure threshold in *Dhdds^R37H+/-^* mice (*P* = 0.03, Welch’s *t* test) but not in WT controls (*P* = 0.345, Welch’s t test)(*n* = 5-6). (**H**) *Dhdds^R37H+/-^* mice show reduced latency to fall in accelerating rotarod test. (**I-J**) AZM (40 mg/kg, i.p., 45-min before the experiment) does not increase latency to fall in the rotarod neither in *Dhdds^R37H+/-^*mice (*P* = 0.42, Welch’s *t* test) nor in WT mice siblings (*P* = 0.87, Welch’s *t* test) (*n* = 7-8). (K-L) *Dhdds^R37H+/-^*mice show reduced discrimination index in the novel object recognition (NOR) test (**K**) and reduced percent of alteration in the Y-maze test (**L**), consistent with impaired short-term and spatial memory, respectively (*n* = 7-13 male and female mice for each genotype). All graphs show individual data, means and SEM. *P*-values were calculated by two-tailed *t*-test with Welch’s correction.

Recently, acetazolamide (AZM), clinically approved for the treatment of glaucoma, epilepsy and intracranial hypertension, was shown to improve titubation, tremor, and generalized edema in an adult patient with autosomal recessive DHDDS-CDG.^36^ Thus, we evaluated the effect of an acute intraperitoneal (i.p.) injection of AZM on PTZ-induced seizures in WT and *Dhdds^R37H+/-^* mice. AZM (40 mg/kg BW, in 8% DMSO in saline) or saline, were administered i.p. 45-min before an intravenous infusion of PTZ until the detection of the first myoclonic seizure and epileptic activity on EEG. AZM did not change seizure susceptibility to PTZ in WT mice (*P* = 0.345, Welch’s *t* test), but it reduced seizure susceptibility by 45% in *Dhdds^R37H+/-^* mice (*P* = 0.03, Welch’s *t* test) to a level comparable to that of WT mice (Fig. 2F-G).

We further characterized the motor and cognitive function in *Dhdds^R37H+/-^*mice compared to WT siblings. The motor coordination in male and female *Dhdds^R37H+/-^*mice was studied by an accelerating rotarod (Fig. 2H). This experiment revealed that, compared to WT siblings, *Dhdds^R37H+/-^* mice had a lower latency to fall suggesting a presence of neuromotor deficits. We, thus, tested whether AZM at the same dose (40 mg/kg BW, IP administration, 45 mins before the experiment) also rescued this phenotype, but found that the drug did not affect the levels of latency to fall in either *Dhdds^R37H+/-^* mice or their WT siblings compared to mice treated with saline only (Fig. 2I-J). Mouse memory and learning were assessed using the Y-maze test, that measures spatial and short-term memory, and the novel object recognition (NOR) test, that measures short-term memory. In the NOR test, *Dhdds^R37H+/-^* mice showed a significant reduction in discrimination index compared to WT siblings, suggesting a short-term memory deficit (Fig. 2K). In a similar fashion, they showed a significantly reduced alternation index in Y-maze test suggesting spatial and short-term memory deficits (Fig. 2L).

### *Dhdds^R37H+/-^* mice show decreased density and maturation of cortical GABAergic interneurons, resulting in reduced cortical inhibitory tone

Brains of *Dhdds^R37H+/-^* mice and their matching WT controls were analyzed at sacrifice to assess for potential pathological changes. There was no difference in the wet weight of brains as well as peripheral organs compared to controls (Supplementary Fig. 3). Immunofluorescence microscopy did not reveal alterations in the brain morphology, density of NeuN^+^ neurons, or the presence of CD68^+^ activated microglia suggestive of microgliosis (Supplementary Fig. 4). In contrast, the density of GFAP^+^ astrocytes was increased in the CA1 area of hippocampus and showed a non-significant trend towards an increase in the cortex, suggestive of astrogliosis (Supplementary Fig. 4). There was, also, a ∼30% decrease in the number of parvalbumin-positive interneurons (PV^+^ INs), a key population of GABAergic INs in the adult brain^37^ (Fig. 3A). Since PV^+^ INs are typically surrounded by the perineuronal nets (PNN), distinctive extracellular matrix structures that stabilize inputs regulating synaptic plasticity and neuronal excitability, we further assessed PNN integrity. For this, brain tissues were labeled with antibodies against PNN-forming proteoglycan, aggrecan, or *Wisteria floribunda* agglutinin (WFA) lectin, specific to chondroitin sulfate chains on PNN. The brains of *Dhdds^R37H+/-^*mice showed >50% reduction in the number of PV^+^ INs surrounded by PNN compared to WT mice (Fig. 3B-C), compatible with INs scarcity or reduced maturity.

**Figure 3.**
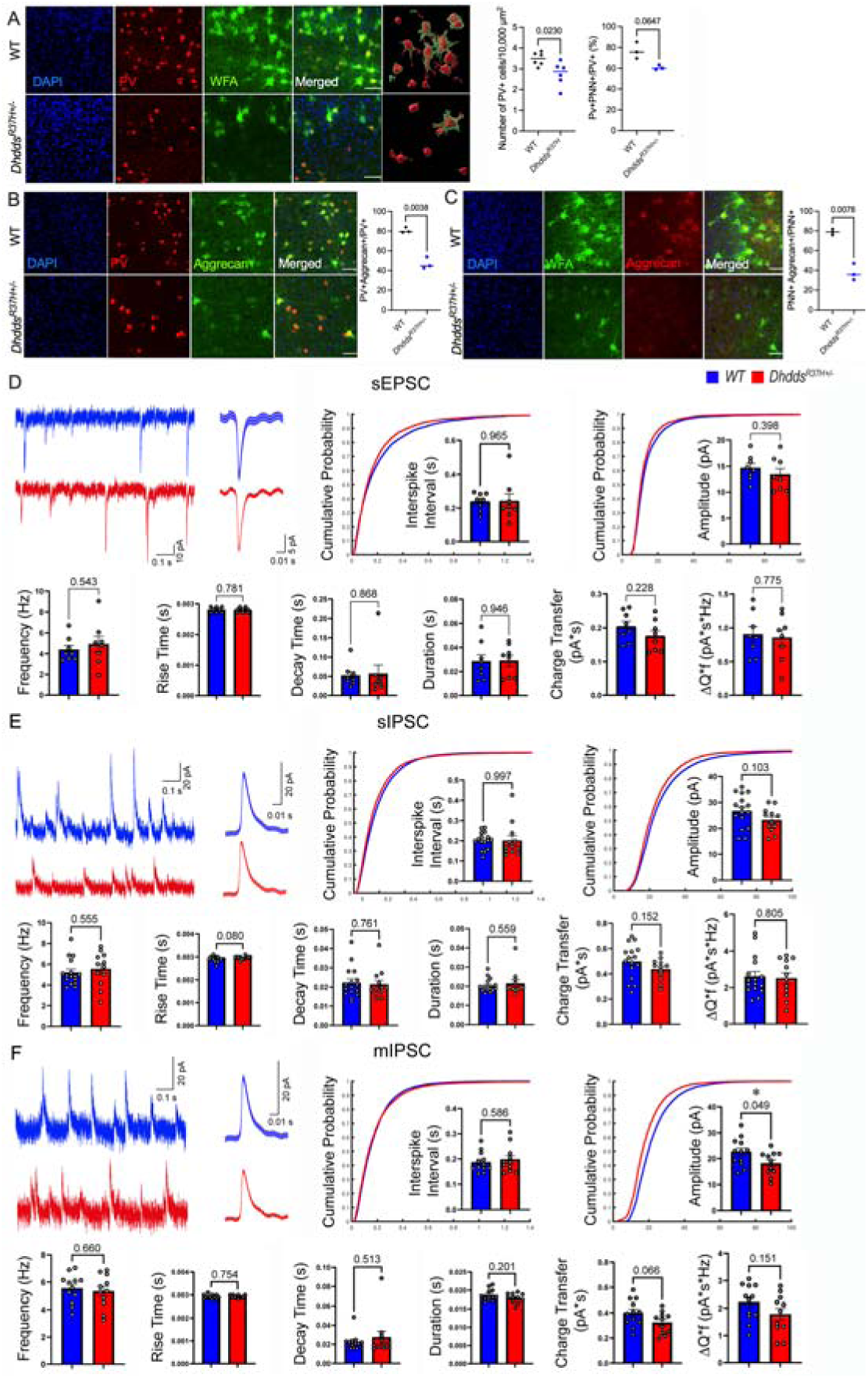
***Dhdds^R37H+/-^*mice show decreased density of cortical GABAergic interneurons of and reduced mIPSC amplitude in layer V pyramidal cells (PCs).** (**A-C**) Representative confocal images and 3D reconstructions of cortical PV interneurons and associated PNNs in the brain of 6-week-old *Dhdds^R37H+/-^*and WT mice. Coronal brain sections were labeled with antibodies against mouse PV (red) and fluorescently labeled WFA lectin (green) or antibodies against aggrecan (green in H, red in I). DAPI (blue) was used as a nuclear counterstain. Size bars are 50 μM. Graphs show quantification of fluorescence by ImageJ software revealing reduction of WFA^+^/PV^+^ and aggrecan^+^/PV^+^ cells in the cortices of *Dhdds^R37H+/-^* compared to WT mice (*n* = 3-6, *P*-values were calculated by unpaired t test with Welch’s correction). (**D**) Representative traces of sEPSCs recorded in PCs from control (blue, n = 8 cells, 5 mice) and *Dhdds^R37H+/-^* (red, *n* = 8 cells, 5 mice) mice with average events (bold trace). Summary bar graphs show no differences for sEPSC frequency, inter-spike interval, event kinetics, amplitude, charge transfer, or ΔQ*f of sEPSCs. (**E**) Representative traces of sIPSCs recorded in PCs from control (blue, *n* = 16 cells, 6 mice) and *Dhdds^R37H+/-^* (red, *n* = 12 cells, 6 mice) mice with average events (bold trace). Summary bar graphs show no differences for sIPSC frequency, inter-spike interval, event kinetics, amplitude, charge transfer, or ΔQ*f of sIPSCs. (**F**) Representative traces of mIPSCs recorded in PCs from control (blue, *n* = 12 cells, 6 mice) and *Dhdds^R37H+/-^* (red, *n* = 11 cells, 5 mice) mice with average events (bold trace). Summary bar graphs show no differences for mIPSC frequency, inter-spike interval, event kinetics, charge transfer, or ΔQ*f. There was a significant decrease in the amplitude (*P* = 0.049, Welch’s t-test) of mIPSCs. Bar graphs represent the mean ± SEM. Cumulative curves show the distribution of all recorded events (WT EPSC, *n* = 5900; *Dhdds^R37H+/-^*EPSC, *n* = 5260; WT IPSC, *n* = 10000; *Dhdds^R37H+/-^*IPSC, *n* = 12515; WT mIPSC, *n* = 8793; *Dhdds^R37H+/-^* mIPSC, *n* = 9975; all events are shown, different numbers of events are recorded from each cell).

To investigate how a loss of INs in the cortex affects local network activity, recordings of excitatory and inhibitory postsynaptic currents on layer V somatosensory cortex (S1) principal cells (PCs) were performed in acute coronal brain slices of P26-35 *Dhdds^R37H+/-^* mice and their WT littermate controls. Whole-cell patch-clamp recordings revealed no change in the frequency, amplitude, decay time, or charge transfer of sEPSCs in layer V PCs (WT: n = 8 cells from 5 mice; *Dhdds^R37H+/-^*: *n* = 8 cells from 5 mice) (Fig.3D). There was also no change in sIPSC parameters (WT: n = 16 cells from 6 mice; *Dhdds^R37H+/-^*: *n* = 12 cells from 6 mice) (Fig.3E). However, there was a significant reduction in mIPSC amplitude (p = 0.049) (WT: *n* = 12 cells from 6 mice; *Dhdds^R37H+/-^*: *n* = 11 cells from 5 mice) in *Dhdds^R37H+/-^* mice (Fig. 3F). These results reveal a maintenance of cortical excitatory currents in acute slices of *Dhdds^R37H+/-^* mice, with a reduction in inhibitory currents received by layer V PCs consistent with a reduction in cortical PV cell maturation/density, as well as potential post-synaptic changes explored below.

### *Dhdds^R37H+/-^* mice reveal changes in protein N-glycosylation

To establish whether *cis-*PTase deficiency and reduced dolichol levels were associated with aberrant or hypoglycosylation of brain proteins, we carried out a glycoproteomic study. The simultaneous MS/MS analysis of glycan and protein sequences of enriched glycopeptides from tryptic digests of total brain homogenates revealed altered levels and structures of glycan chains on multiple brain glycoproteins, with most glycopeptides downregulated (Fig. 4A). This reduction extended to the peptides bearing both oligomannose and complex glycans (Fig. 4B). Further, detailed analysis of glycopeptides with altered structures revealed a drastic decrease in those with mature forms of glycans and GlcNAc bisected glycans and simultaneous increase in those with smaller/truncated glycans (Fig. 4C-D). Similar truncated species were previously reported in plasma of CDG I (ALG1, PMM2, MPI and ALG13) patients.^38,39^ The mostly reduced glycopeptides included those involved in neuronal development and function, such as Neural cell adhesion molecule 1 (NCAM1), Neural cell adhesion molecule L1-like protein (NCHL1), Embigin (EMB), Protocadherin (FAT2), and Contactin-4 (CNTN4) or synaptic transmission, such as cell adhesion molecule 3 (CADM3), Neuroplastin (NPTN), Synaptic vesicle glycoprotein 2B (SV2B) or glutamate receptor 2 (GRM2) (Fig. 4C-D). Also of interest was a reduction of the major prion protein (PRNP), involved in neuronal development, synaptic plasticity and neuronal myelin sheath maintenance, GDNF family receptor alpha-2 (GFRA2), the receptor for neurturin that supports neuronal survival^40–43^, and A-type potassium channel modulatory protein (DPP6), which modulates activity of the major neuronal potassium channel KCND2^44^ (Fig. 4C-D).

**Figure 4.**
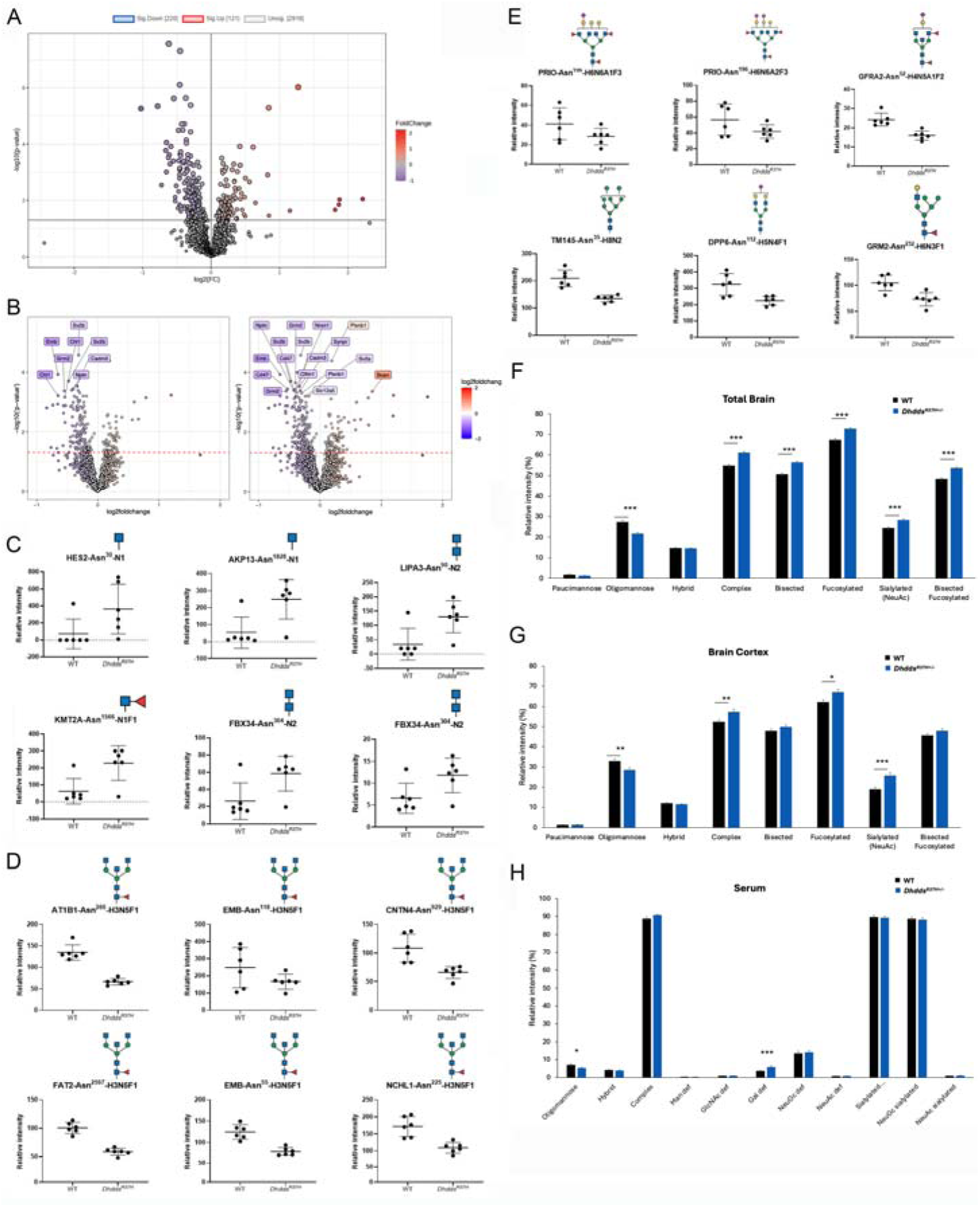
Aberrant N-glycosylation in *Dhdds^R37H+/-^* mouse tissues. (**A**) Glycoproteome analysis reveals global reduction of N-linked glycopeptides in the *Dhdds^R37H+/-^* mouse brain. (**B**) Glycopeptides carrying both oligomannose and complex types of glycans are downregulated in *Dhdds^R37H+/-^*mouse brain. Six most increased (**C**) and most decreased (**D)** glycopeptides in *Dhdds^R37H+/-^* mouse brain. (**E**) Other glycopeptides of interest decreased in *Dhdds^R37H+/-^* mouse brain (*n* = 6, 3 males and 3 females for each genotype). (**F-H**) Levels of N-glycans are altered in *Dhdds^R37H+/-^* mice. Graphs show relative abundance of specific protein N-glycan populations in total brain (**F**), brain cortex (**G**) and serum (**H**) of *Dhdds^R37H+/-^* mice and their WT siblings. Data are expressed as means (+/- SEM) of the peak areas normalized to the total areas of individual species grouped as shown in Tables S1A (brain and cortex) or S1B (serum). Statistical analyses were performed using one-way ANOVA; * *P*< 0.05, ** < 0.01 and *** < 0.001 (*n* = 4, 2 males and 2 females for each genotype).

To confirm these results, we analyzed the N-glycome profile of the whole brain, brain cortex and serum of WT and *Dhdds^R37H+/-^* mice by MALDI-TOF MS and MS/MS (Fig. 4F-H, Supplementary Fig. 5). N-glycan structures were assigned based on the MS and MS/MS spectra by comparing them with previously published data.^45^ In the *Dhdds^R37H+/-^* whole brain samples, we observed a reduction in oligomannose-containing N-glycans coinciding with drastic increases in all complex glycan species, either bisected or fucosylated/sialylated (most of them with terminal Neu5Ac) compared to WT mice (Fig. 4F). The brain cortex followed the same trend (Fig. 4G). Changes in N-glycans of serum proteins were less pronounced. Nevertheless, a reduction in oligomannose structures was detected together with an increase in truncated undergalactosylated complex N-glycans, suggesting a galactosylation defect for serum glycoproteins in *Dhdds^R37H+/-^* mice (Fig. 4H). Together, these results establish for the first time that the *DHDDS* p.R37H variant causes a dual CDG (combined type I and type II defect) and that it is associated with a distinct abnormal glycosylation of brain and serum proteins.

### Dysregulated gene expression profiles and proteome alterations in the CNS of *Dhdds^R37H+/-^* mice

To get further insights into the pathological changes in the CNS of *Dhdds^R37H+/-^* mice, we analysed gene expression by total RNA sequencing and protein abundance by semi-quantitative isotope-free proteomics. RNA sequencing revealed a drastic reduction in the expression of genes associated with GABAergic synaptic transmission, synapse assembly, CNS development, and anterograde trans-synaptic signaling (Fig. 5A). Of special interest was a reduced expression of the *Hap1* gene encoding for Huntingtin-associated protein 1 involved in delivery of GABA receptors to synapses and their recycling, the *Gabr2* gene encoding for the GABA receptor subunit 2, *Syn2* gene encoding for Synapsin 2, and *Bsn,* encoding for the GABAergic presynaptic scaffold protein Bassoon (Fig. 5B). We also detected a major reduction in the expression of *Cacna1a, Cacna1b, Cacna1i,* and *Cacna1h* genes that encode subunits of Voltage-dependent P/Q-type and N-type calcium channels pivotal in neuronal function (Fig. 5B). Pathogenic variants in the *Cacna1a* and *Cacna1b* genes cause Spinocerebellar ataxia 6 (SCA6) and Neurodevelopmental disorder with seizures and non-epileptic hyperkinetic movements (NEDNEH) with the symptoms resembling those in DHDDS-CDG.^46^ We further studied CACNA1A protein levels in the mouse brain homogenates by immunoblotting (Supplementary Fig. 6) and confirmed that in the *Dhdds^R37H+/-^* mouse brains, the protein is drastically reduced compared to the WT siblings. RNA sequencing also revealed downregulation of genes associated with cholesterol biosynthetic and metabolic processes, as well as steroid and isoprenoid metabolic processes, while expression of genes associated with the glycosylation pathways (GalNAc and GlcNAc transferase, galactosyltransferase, sialyltransferase, and others) was increased, potentially in attempt to compensate for low dolichol levels (Fig. 5A).

**Figure 5.**
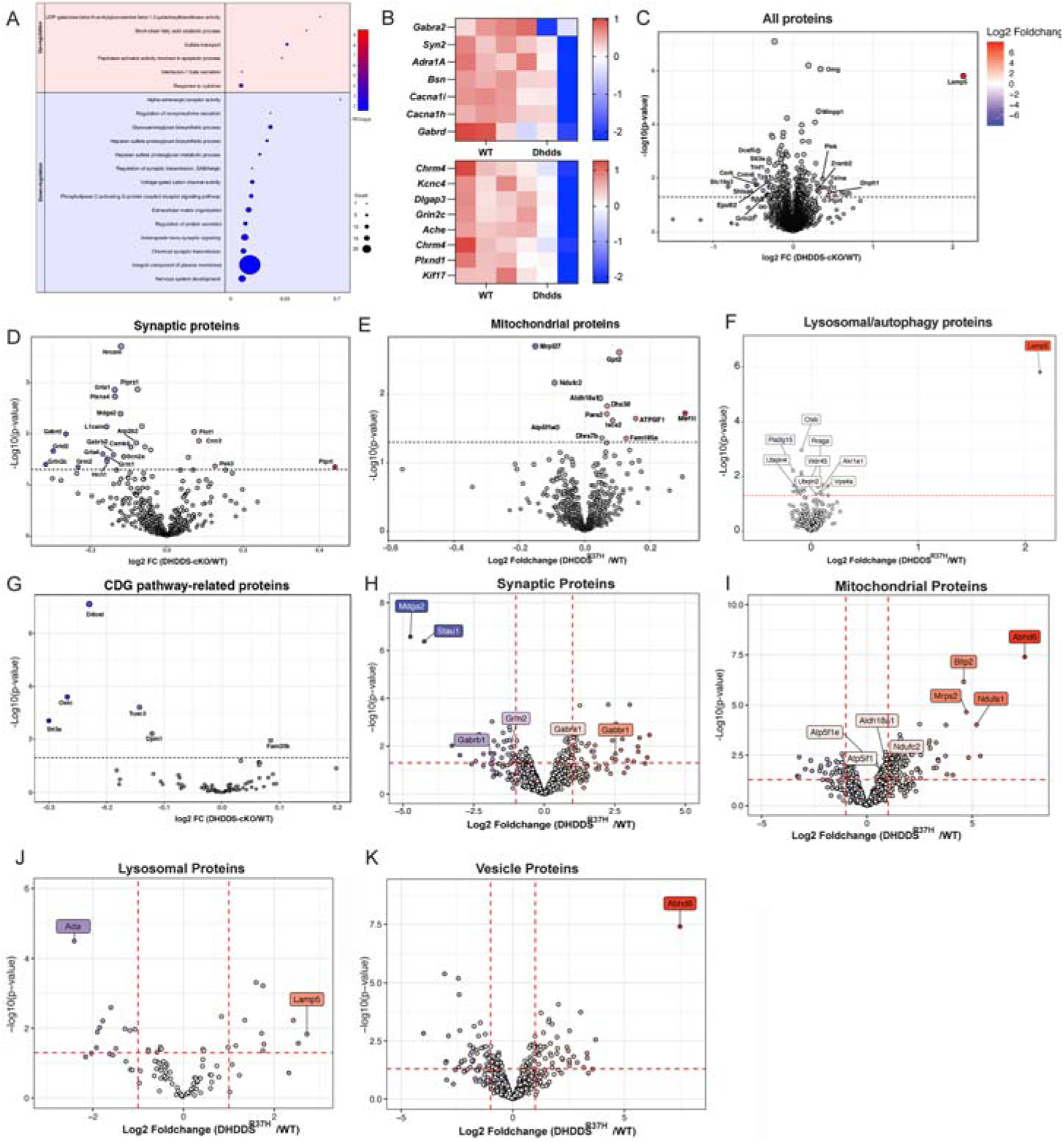
Multiomics analysis unravels CNS pathophysiology in *Dhdds^R37H+/-^* mice. (**A-B**) RNA sequencing analysis of brain tissues reveals a drastic reduction in the expression of genes associated with synaptic transmission, nervous system development, anterograde trans-synaptic signaling and glycosaminoglycan (GAG) synthesis pathways. (**A**) Dot plots show significantly enriched GO terms for the genes upregulated and downregulated in the hippocampi of *Dhdds^R37H+/-^*compared to WT mice. (**B**) Heat maps of the genes of interest significantly upregulated and downregulated in *Dhdds^R37H+/-^* compared to WT mice. GO terms are plotted in the order of gene ratios, and each pathway is shown as a circle with the color representing the *p*-values (−log10) and the size representing the number of differentially expressed genes. The heatmap colors and their intensity show changes in gene expression levels. Data were obtained by sequencing mRNA samples extracted from 3 mice per genotype. (**C**) Global alterations in the brain proteome of *Dhdds^R37H+/-^* compared to WT mice. Alterations in synaptic (**D**), mitochondrial (**E**) and lysosomal/autophagy (**F**) proteins in brains of *Dhdds^R37H+/-^* mice. (**G**) Alterations in the levels of proteins of glycosylation machinery and/or CDG pathway-related proteins in the brains of *Dhdds^R37H+/-^* mice (*n* = 6, 3 males and 3 females for each genotype). (**H-K**) Semiquantitative proteomic analysis of brain synaptosomes confirms alterations in the levels of synaptic proteins (**H**), mitochondrial proteins (**I**), lysosomal proteins (**J**), and proteins involved in vesicular transport (**K**).

Notably, we also detected a massive reduction in the expression of genes associated with glycosaminoglycan (GAG) biosynthesis pathways, including heparan sulfate (HS) synthetic processes, proteoglycan biosynthetic process, HS sulfotransferase activity and HS proteoglycan metabolic process (Fig. 5A). This included the aggrecan proteoglycan (*Acan*) gene consistent with the decrease in the number of aggrecan-containing PNNs in the mouse brain (Fig. 3B-C). To test whether downregulation of GAG synthesis genes was associated with low brain levels of GAGs, we analyzed their levels by enzyme digestion followed by mass spectrometry. Heparan sulfate, keratan sulfate, and chondroitin sulfate levels were similar in the brains of *Dhdds^R37H+/-^* and WT siblings but increased in the tibia bone of *Dhdds^R37H+/-^* mice (Supplementary Fig. 7).

Semi-quantitative proteomic analysis revealed drastic alterations in the CNS proteome of *Dhdds^R37H+/-^* mice (Fig. 5C) with a significant downregulation of proteins associated with ion transport, protein N-linked glycosylation, synaptic transmission, axon guidance and neuron projection development pathways (Supplementary Fig. 8). In turn, the cellular component analysis revealed major alterations in the levels of synaptic (Fig. 5D), mitochondrial (Fig. 4E), and lysosomal/autophagic (Fig. 5F) proteins. Most of the synaptic proteins were reduced, including neuronal cell adhesion proteins NRCAM and L1CAM, subunits of glutamate receptors (GRIA1, GRIA4, GRLN2C, GRID2, GRM1, and GRM2), subunits of GABA receptors (GABRD and GABRB2) and potassium/sodium cyclin gated channel HCN1 and Ca^2+^ transporting ATPase, ATB2. Of special interest was a major decrease in the glutamate signalling-coupled sodium channel, SCN2A, which genetic deficiency causes Benign familial neonatal-infantile seizures, and the inhibitory synapse organizer MDGA2 (Fig. 5D). Both mitochondrial and lysosomal/autophagic proteins showed a generally dysregulated patterns with some proteins decreased and others increased (Fig. 5E-F). Such changes are generally consistent with partial mitochondrial dysfunction and blocks in autophagy and lysosomal catabolism.

Finally, we detected alterations in glycosylation machinery proteins and/or those previously associated with other CDG (Fig. 5H). In particular, levels of oligosaccharyltransferase complex subunit, OSTC, Dolichyl-diphosphooligosaccharide-protein glycosyltransferase subunit TUSC3, Dolichyl-diphosphooligosaccharide-protein glycosyltransferase, DDOST, Dolichol-phosphate mannosyltransferase subunit 1, DPM1, and Dolichyl-diphosphooligosaccharide-protein glycosyltransferase subunit, STT3A, were decreased. Genetic deficiencies of the last 3 proteins are causative for DDOST-CDG, STT3A-CDG and DPM1-CDG CDG1R, CDG1E and CDG1W, while pathogenic variants in the *TUSC3* interfering with N-glycosylation of Cys proximal acceptor sites cause intellectual developmental disorder, autosomal recessive 7 (MRT7), also described as TUSC3-CDG.^3,47^ The levels of enzymes involved in cholesterol biosynthesis and transport were not altered.

To confirm the alterations in the levels of synaptic proteins, we conducted a proteomic analysis of purified mouse brain synaptosomes.^48^ Synaptosomes were isolated from *Dhdds^R37H+/-^*and WT mice by differential centrifugation, and their protein content was analyzed by label-free LC-MS/MS. Based on immunoblot analysis, the synaptosome preparation was >3-fold enriched in synaptic proteins compared to the whole brain homogenate. In total, levels of 2957 proteins were compared between the synaptosomes from WT and *Dhdds^R37H+/-^* mice, and proteins with significantly altered levels were classified according to their biological function and linked to a particular metabolic or signaling pathway using automated gene ontology annotation.^49^ The synaptic and lysosomal proteins, as well as those involved in vesicle trafficking were downregulated in *Dhdds^R37H+/-^* synaptosomes, whereas mitochondrial proteins, increased, consistent with the results of total brain proteome analysis (Fig. 5H-K, Supplementary Fig. 8). Specifically, the reduction of MDGA2 protein, GABA receptor subunits, glutamate receptor 2 subunits, and Dolichyl-diphosphooligosaccharide-protein glycosyltransferase subunits, identified by whole brain proteomics, was confirmed by the analysis of synaptosomes.

### Brains of *Dhdds^R37H+/-^* mice show abnormal lipid composition

Since dolichol is an essential component of cellular membranes and is involved in the pathways related to lipid biosynthesis, we conducted a non-targeted lipidomic analysis of the mouse brain tissues by LC-MS/MS. This experiment revealed an increase in several species of phosphatidylinositol (Fig. 6A), phosphatidylserine, (Fig. 6B), and phosphatidylcholine (Fig. 6C) together with a decrease in phosphatidic acid (Fig. 6D) and triacylglycerols (Fig. 5E) in *Dhdds^R37H+/-^* mice. Notably, our proteomic analyses revealed changes in levels of multiple enzymes involved in processing or hydrolysis of these lipid species. In particular, 1-phosphatidylinositol phosphoinositide phospholipase C-gamma-2 (PLCG2), Calcium-independent phospholipase A2-gamma (PNPLA8) and Phospholipase A-2-activating protein (PLAA) were decreased in the synaptosomes of *Dhdds^R37H+/-^*mice (Fig. 6F). On the other hand, monoacylglycerol lipase ABHD6 that preferentially hydrolyses medium-chain saturated monoacylglycerols was drastically increased. Additionally, proteomics analysis of whole brain tissues revealed an increase in the levels of PLD3, a transphosphatidylase which catalyzes the R to S stereo-inversion of the glycerol moiety between (S,R)-lysophosphatidylglycerol and monoacylglycerol substrates, the lysosomal phospholipase A PLA2G15, diacylglycerol lipase-alpha (DAGLA), as well as putative lipases PLBD2 and GDPD5 (Fig. 6G).

**Figure 6.**
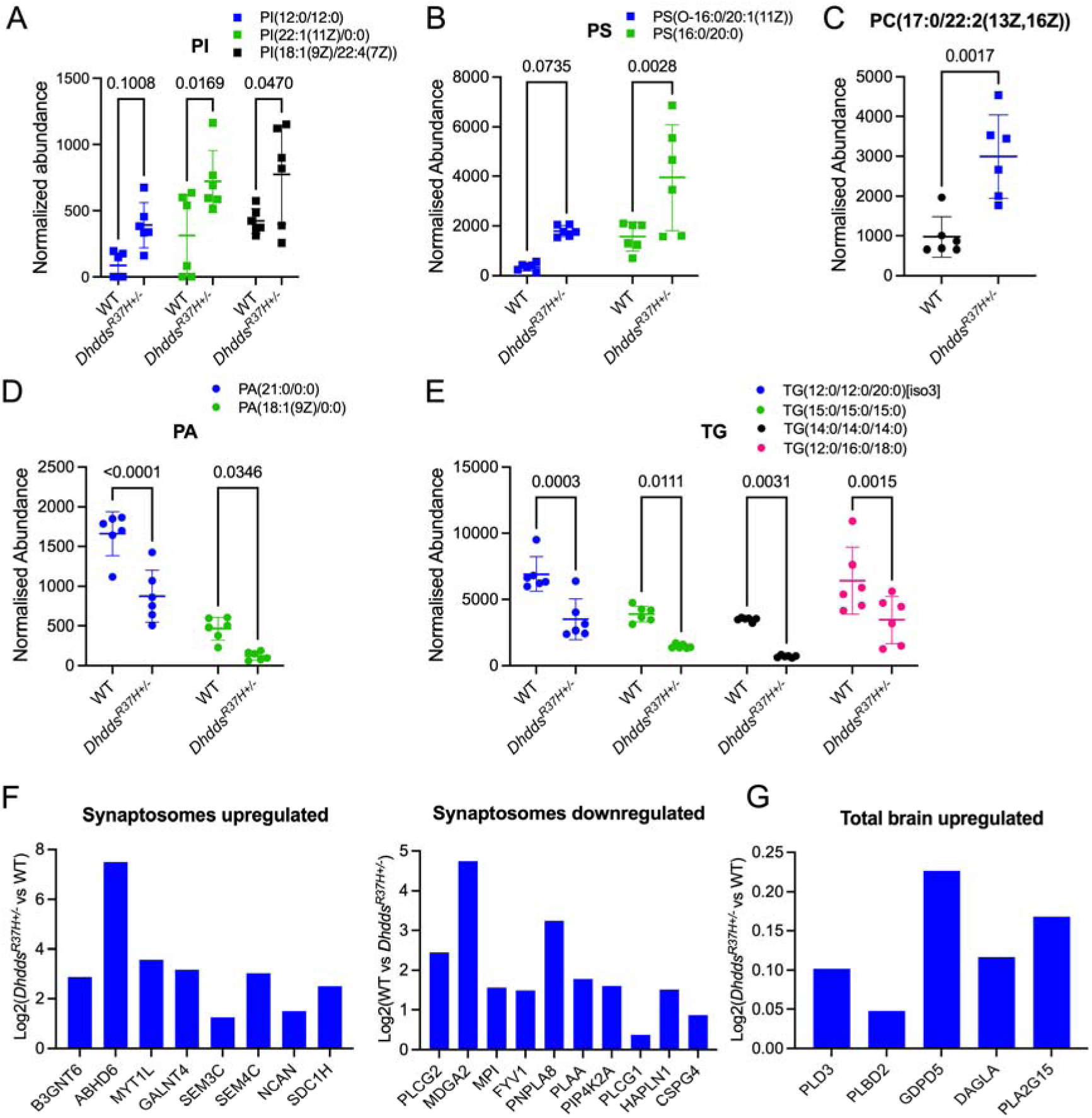
Abnormal phospholipid and triglyceride composition and alterations in the lipase levels in the brains of *Dhdds^R37H^*^+/-^ mice. (**A-E**) Untargeted lipidomic analysis reveals increased levels of phosphatidylinositol **(A)**, phosphatidylserine **(B)**, phosphatidylcholine **(C)**, and decreased levels of phosphatidic acid **(D)** and triglycerides **(E)** species (*n*=6, 3 males and 3 females for each genotype, *p* values were calculated by unpaired *t* test). (**F-G**) Altered levels of phospholipid and triglyceride lipases in the synaptosomes (**F**) and total brains (**G**) of *Dhdds^R37H+/-^* mice.

As alterations in the levels and/or distribution of glycosphingolipids (GSL) were previously reported in the cultured skin fibroblasts of PME patients with DHDDS pathogenic variants,^50^ we assessed GSL brain levels by targeted analysis. After extraction and partial purification, fluorescently labelled GSL glycans were analysed by normal phase HPLC. This analysis did not reveal differences in the GSL content and composition between *Dhdds^R37H+/-^* and WT mice (Supplementary Fig. 9A-E). Neither did we detect changes in the levels of several GSL species identified independently by LC-MS/MS analysis (Supplementary Fig. 9F-H) or MALDI MSI imaging (not shown). Both techniques, however, revealed a drastic reduction of sulfatides, most pronounced in the cerebellum, medulla and mid-brain regions (Fig. 7).

**Figure 7.**
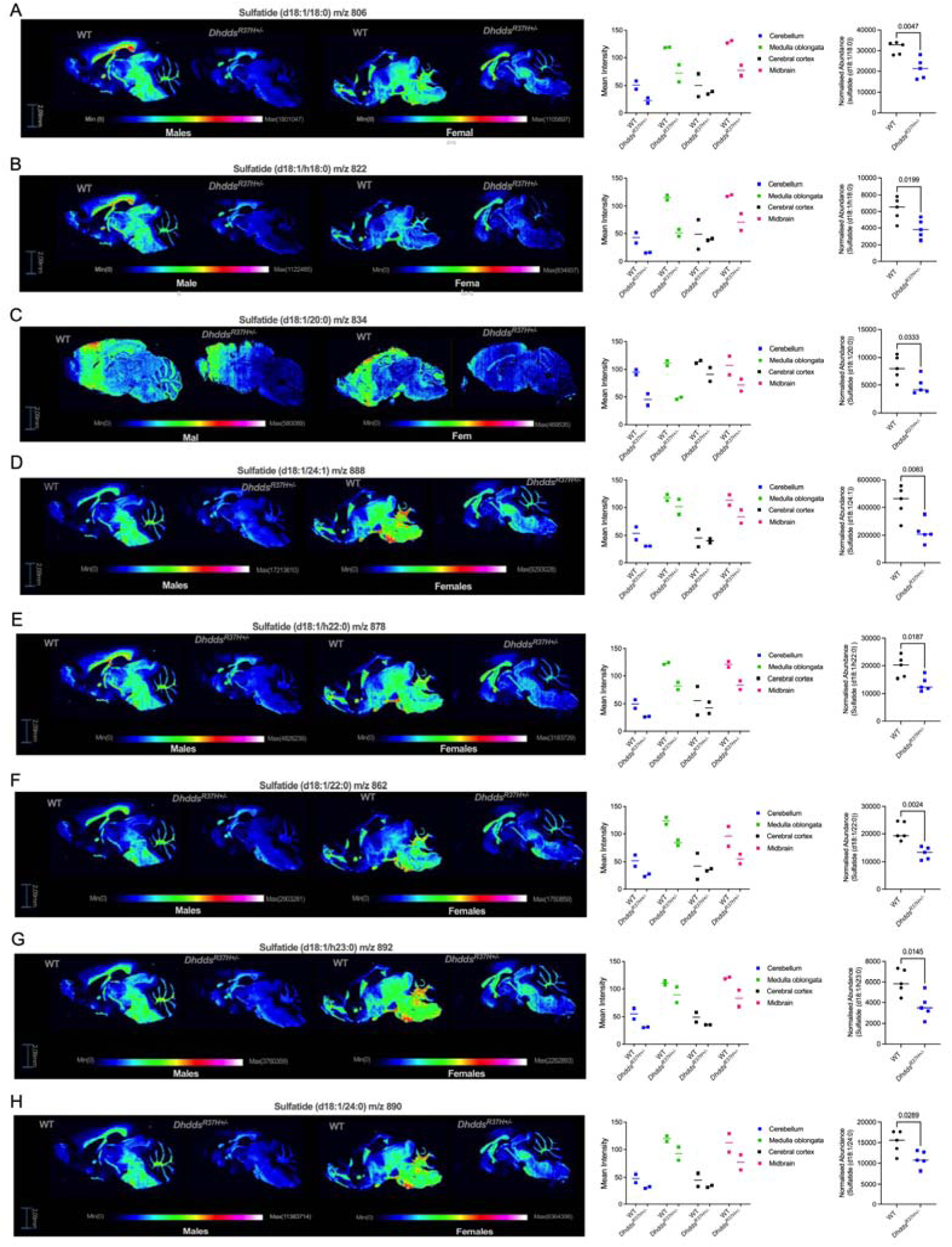
MALDI-MSI and LC-MS/MS analyses depicting reduced levels of sulfatides in the brains of *Dhdds^R37H^*^+/-^ mice. Spatial distribution and levels of sulfatide (d18:1/18:0) (**A**), sulfatide (d18:1/h18:0) (**B**), sulfatide (d18:1/20:0) (**C**), sulfatide (d18:1/24:1) (**D**), sulfatide (d18:1/h22:0) (**E**), sulfatide (d18:1/22:0) (**F**), sulfatide (d18:1/h23:0) (**G**), **and** sulfatide (d18:1/24:0) (**H**) were estimated by MALDI-MSI (*n* = 2, one male and one female in each group, left and central panels) and untargeted lipidomic LC-MS/MS analysis in isopropanol/acetonitrile/water (2:1:1) extracts of brain tissues of *Dhdds^R37H+/-^* and WT mice (*n* = 5, 3 males and 2 females per genotype, right panels). All graphs show individual results, means and SD. *P*-values were calculated by *t*-test.

**Figure 8.**
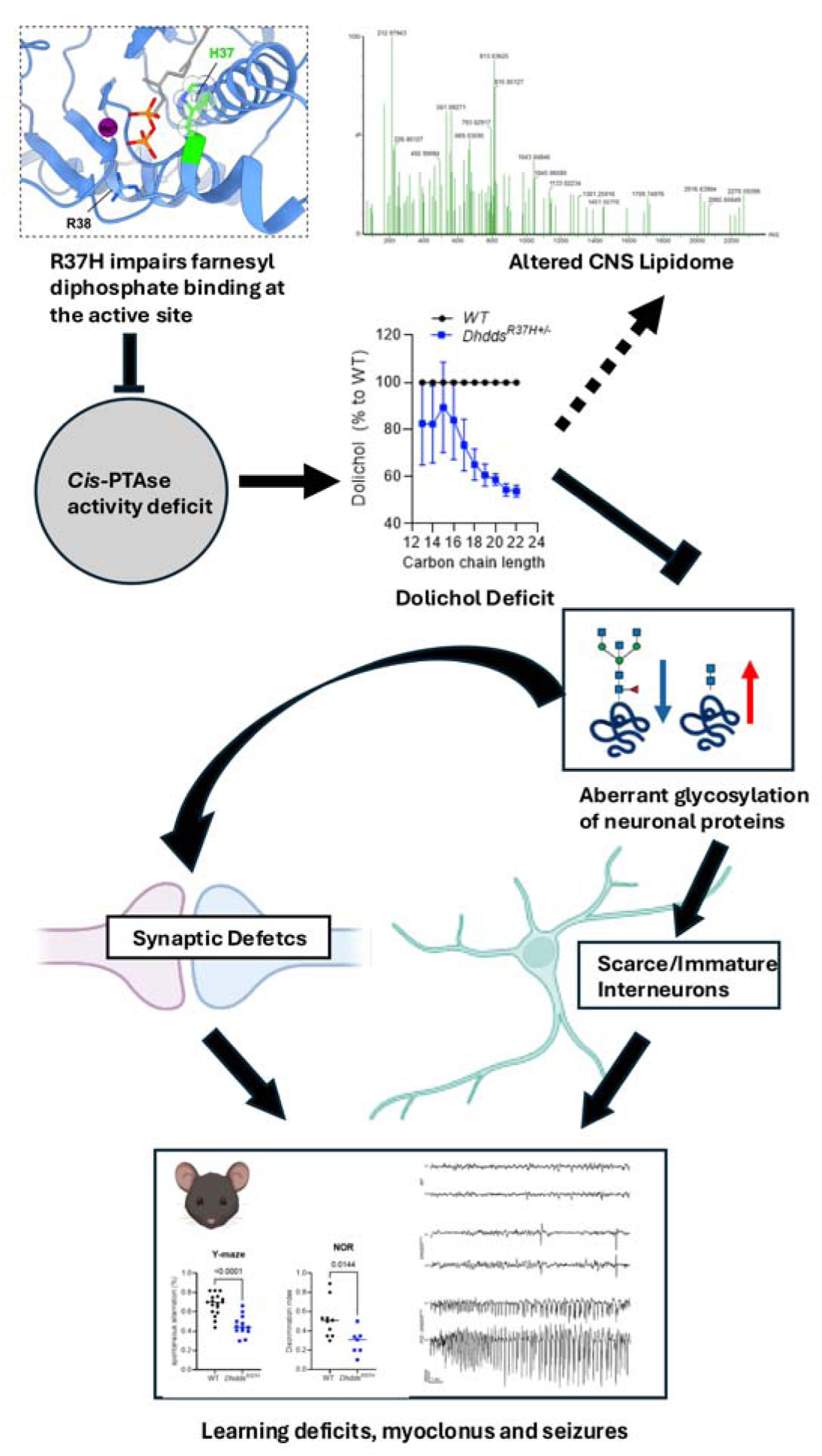
Putative biochemical mechanism underlying the pathological CNS phenotype in *Dhdds^R37H^*^+/-^ mice. The R37H DHDDS variant impairs farnesyl diphosphate binding at the active site and abolishes the catalytic activity of *cis*-PTAse. These results in ∼50% reduction of the enzyme activity in the brain tissues, deficient levels of dolichol and altered spectra of dolichol species. As a result, multiple brain glycoproteins including those involved in synaptic transmission show hypoglycosylation and/or aberrant structure of N-linked glycan species leading to GABAergic synaptic defects, immaturity/scarcity of cortical interneurons, excitation/inhibition (E/I) imbalance, seizures, memory deficits and neuromuscular defects. Major perturbations in the CNS proteome and lipidome may be a consequence of dolichol deficit or the secondary changes induced by seizures.

## Discussion

Recent advances in genetic testing and screening have revealed a dramatic increase in the number of diagnosed patients affected with *DHDDS/NUS1* pathogenic variants. However, effective specific treatments are still lacking. Moreover, our understanding of the mechanism of the disease remains very limited, emphasizing the need for additional studies, including those involving animal models. In the current study, we generated a mouse strain with the recurrent human pathogenic DHDDS variant p.R37H associated with a severe clinical phenotype, that included a developmental delay, cognitive deficits, progressive myoclonus, tremor, ataxia and seizures.^7^

In homozygosity, the p.R37H variant resulted in early embryonic lethality, while heterozygous mice were fertile and had a normal survival. They, however, showed unfrequented interictal spiking activity (on average one episode/h), as well as spontaneous seizures. Mice also revealed neuromuscular and memory deficits. Overall, these results suggest that the model has strong face validity, as the mouse phenotype was similar to that of human patients. Notably, the seizure threshold in mice was restored by acute treatment with AZM, a carbonic anhydrase inhibitor clinically approved for the treatment of glaucoma, epilepsy, and intracranial hypertension. The drug accelerates the elimination of bicarbonates and other electrolytes, resulting in an overall reduction of osmotic pressure. Previously, we used the drug on a compassionate basis to treat an adult female patient homozygous for the DHDDS K42E variant and presenting with seizures, ataxia, protein-losing enteropathy, tremors, and titubation in association with a retinitis pigmentosa. AZM improved titubation, tremor, and generalized edema, although, the patient still had mild ataxia.^36^ The effect of the drug on seizures was not evaluated as they have been well controlled with levetiracetam.^36^ Our current results provide further support for a potential therapeutic benefit of AZM in reducing seizure susceptibility. Previously, it was hypothesized that AZM reduces seizures in CDG by inhibiting hypoglycosylated and overactive calcium channel CACNA1A through lowering the intracellular pH and reducing the excitability of neuronal membranes.^51^ This mechanism is consistent with our glycoproteomics results indicating that CACNA1A in *Dhdds^R37H+/-^* mice shows a reduced glycosylation (log2 fold change -0.21; *P*-value 0.04). AZM had been previously shown to increase PTZ seizure threshold in WT mice, although at much higher doses. In particular, 82.7 mg/kg BW dose of AZM suppressed 50% of seizures induced by subcutaneous PTZ in WT mice.^52^ In the current study, an AZM dose of 40 mg/kg BW was sufficient to increase the seizure threshold to i.v. PTZ in *Dhdds^R37H+/-^*mice but not in WT mice, showing a higher sensitivity to the antiseizure properties of AZM in our model. On the other hand, AZM at this dose did not show efficacy in rescuing neuromuscular deficits in the accelerating rotarod test.

Our structural modeling and the analyses of enzymatic activity of the recombinant mutant enzyme indicated that the R37H variant abolishes the catalytic function, consistent with ∼50% reduced *cis*-PTase activity in the brain tissues of the *Dhdds^R37H+/-^* mice. This level of activity is not sufficient to sustain normal levels of dolichol production, as demonstrated by a strong reduction of polyprenols and dolichols in the brain of *Dhdds^R37+/-^* mice, reconfirming that *cis-*TPase is the rate-limiting enzyme of the dolichol pathway. Interestingly, we also observed a change in the spectrum of dolichol species, with the most profound decrease observed for the dolichols with longer carbon chains (C19-C22) and suggesting alteration in the enzyme specificity. Of note, similar observations were made in human patients harboring the retinitis pigmentosa-associated DHDDS variant K42E and the CDG-associated NgBR variant R290H.^53,54^ The increase in shorter products supports the previously proposed “tug-of-war” mechanism for polyprenol synthesis, where product release is dictated by the opposing forces of interactions within the active site or the ER membrane.^18^ The contribution of this change in dolichol spectrum to the observed biological and clinical phenotype merits future studies.

Glycoproteomic analysis of brain tissues and the analyses of N-linked glycans in the brain and serum demonstrated that impaired dolichol synthesis results in severe hypo-N-glycosylation. It also alters N-glycosylation of multiple proteins by drastically reducing the levels of those bearing oligomannose-containing and mature glycan species. These changes resemble those detected in CDG type I disorders, typically caused by deficiencies of the enzymes of the oligosaccharyltransferase (OST) complex affecting the GlcNAc2Man9Glc3 glycan assembly and transfer from the dolichol phosphate to the N-glycosylation site(s) of the recipient proteins. Interestingly, several glycosylation machinery proteins, which genetic defects were previously associated with other CDG type I, were reduced in *Dhdds^R37H+/-^*mice. In particular, we detected reduced levels of STT3A and DDOST, both components of the OST complex, catalyzing the transfer of oligomannose oligosaccharides to asparagine residues on nascent polypeptides in the ER. DPM1, dolichol-phosphate mannosyltransferase subunit, which synthesizes Dol-P-Man from GDP-mannose and dolichol-phosphate, was also deficient. Further studies should clarify if and how these changes can contribute to protein hypoglycosylation in *Dhdds^R37H+/-^* mice. In addition, N-glycome analysis revealed a change in the levels of glycan species. Oligomannose structures were reduced while complex, bisected, fucosylated and sialylated glycans, increased in the brains of *Dhdds^R37H+/-^* mice. Such changes are characteristic for CDG type II disorders and caused by the genetic defects of enzymes catalysing the glycan processing steps that occur after the GlcNAc2Man9Glc3 transfer. Thus, likewise many other CDG I subtypes,^55,56^ DHDDS-CDG can be described as a CDG type I/type II disorder involving, both, overall reduction of N-glycosylation sites, resulting from dolichol biosynthesis defect, and alterations in the structure of the N-linked glycan species likely associated with secondary changes in the expression of glycan-processing enzymes. Interestingly, no significant changes were observed in the expression of the genes involved in the endoplasmic reticulum-associated protein degradation (ERAD) pathway, suggesting that glycosylation defects did not lead to major misfolding of proteins and indicating that the overall impact of the R37H variant might be limited, yet functionally essential.

Notably, not only the levels of N-glycans or glycopeptides were changed in the brains of *Dhdds^R37H+/-^* mice, but also the levels of the proteins themselves were drastically altered. Both decreased and increased levels were observed for proteins involved in the mitochondrial energy production and autophagy/lysosomal degradation, although most of the synaptic proteins or those engaged in protein glycosylation were strongly decreased. These changes could be secondary to aberrant glycosylation of the proteins in question, known to be essential for their stability and/or proper intracellular trafficking. They also could originate from dysregulated expression of the corresponding genes.

RNA sequencing and proteomics revealed major downregulation (>30) of genes and proteins involved in synaptic transmission (especially those of the GABAergic synapses). This included GABA receptor subunits and the cerebrospinal ataxia causative gene *Cacna1a,* whose protein product, the Voltage-dependent P/Q-type Cav2.1 channel alpha subunit was also hypoglycosylated. Genes associated with the GAG biosynthesis pathway, including the *Acan* gene encoding for the PNN-forming proteoglycan, aggrecan, were also downregulated. Immunofluorescence microscopy confirmed deficits in both aggrecan protein and chondroitin sulfate chains of the PNNs on cortical INs of *Dhdds^R37H+/-^*mice. The analysis also revealed a reduction in numbers of PV^+^ and WFA^+^/aggrecan^+^ cells in *Dhdds^R37H^* mice, consistent with either reduced production or increased apoptosis of PV+ interneurons. Alternatively, INs fated to become PV^+^ neurons may still be present but remain immature (showing no or low PV expression and less developed PNN labeled with WFA and aggrecan). Other IN types or the whole GABAergic neuron population density were not analysed, so we cannot discriminate between these two hypotheses. However, both outcomes would hinder synaptic plasticity and disrupt the excitatory/inhibitory balance within the neural circuits, contributing to the observed epilepsy and cognitive impairments.^37^ This was directly supported by voltage-clamp recordings of network activity in the mouse primary sensory cortex (S1). These experiments revealed a reduction in the amplitude of miniature inhibitory currents received by layer V PCs, consistent with a reduction in cortical PV cell maturation, and post-synaptic changes at GABAergic synapses. Notably, the overall levels of glycosaminoglycans were not changed in the brains but increased in the bone tissue of *Dhdds^R37H+/-^*mice. These alterations may partially explain facial dysmorphisms observed in some DHDDS-CDG patients^12^.

When interpreting changes in the levels of RNA species and metabolites, it is essential to bear in mind that the spontaneous seizures observed in *Dhdds^R37H+/-^* mice can, by themselves, result in massive changes in multiple biological systems. Some of these changes can contribute to the phenotype (being either compensatory or causative) but other can be bystander. Thus, the biological significance of the systemic alterations detected by multiomics analysis would need to be verified experimentally in the future studies.

The untargeted lipidomic analysis of the brain tissues revealed increase in two forms of cholesterol, 7-dehydrocholesterol and dihydrolathosterol. Concentrations of cholesterol and other sterol species were either similar for *Dhdds^R37H+/-^* and WT siblings or showed a non-significant trend for an increase. These results suggest that heterozygosity for the *Dhdds^R37H^* allele drastically reduces dolichol synthesis but does not cause major alterations in branching pre-squalene or post-squalene parts of the mevalonate pathway. As increased cholesterol accumulation has been previously reported in the patient’s fibroblasts,^14^ it remains to be studied whether an increase in cholesterol levels can be observed in other mouse tissues or at a later stage of the disease progression.

Somewhat unexpectedly, lipidomic analysis also revealed changes in triglyceride and phospholipid profiles with a two-fold reduction in triglyceride species and an increase in phosphatidylinositol, phosphatidylserine and phosphatidylcholine together with a decrease in their precursor, phosphatidic acid, suggesting a novel link between the lipid metabolism and protein glycosylation. Also, of interest is a reduction of an essential myelin component, sulfatide in the brains of *Dhdds^R37H+/-^*mice, specifically pronounced in cerebellum, the brain region responsible for coordinating movement, maintaining balance, and controlling motor functions affected in our animal model. Recently, a loss of brain sulfatide was described in Alzheimer disease (AD) cases and animal models and associated with neuroinflammation^57^. Further studies are required to evaluate the impact of lipid alterations in the *Dhdds^R37H+/-^*mice, but, independently of the mechanism underlying changes in the lipid profiles, our results demonstrate that the disruption of dolichol synthesis in the brains of *Dhdds^R37H+/-^* mice not only impacts glycoprotein biosynthesis but also affects composition and, perhaps, structural properties of cell membranes. This could have cascading effects on neuronal function, contributing to the cognitive deficits revealed by the behavioral analyses.

One question which remains unanswered by this study is whether the phenotypic changes in *Dhdds^R37H+/-^* mice and, potentially, in patients heterozygous for pathogenic missense *DHDDS* variants, are mainly caused by the haploinsufficiency (a ∼50% reduction of *cis*-PTase activity and dolichol levels), or by a gain-of-function/dominant negative effect of the mutant DHDDS protein. This question is essential for future selection of therapeutic strategies either based on gene replacement therapy in the case of pure loss of function or on ASO-mediated inhibition of mutant DHDDS expression. Considering that *cis*-PTase has a heterotetrametric structure containing two DHDDS and two NUS1 proteins, we cannot exclude formation of the enzyme complex containing one mutant and one WT DHDDS protein, where the mutant subunit can induce a conformational change potentially affecting the activity of the WT variant. However, the comprehensive answer to this question can be obtained only by comparing the phenotypes of *Dhdds^R37H+/-^* and *Dhdds^+/-^* heterozygous knockout mice which is currently in progress in our laboratory.

In summary, our findings provide a framework for understanding the underlying mechanisms of genetic diseases associated with pathogenic *DHDDS* variants. They also demonstrate that the *Dhdds^R37H^* strain is a valid disease model, potentially suitable to explore targeted therapeutic strategies, such as gene therapy or mutation repair, aimed at restoring dolichol levels to ameliorate the observed synaptic defects and metabolic disturbances. We showed for the first time that monoallelic pathogenic variants in *DHDDS* cause a novel type I/II congenital disorder of glycosylation. Our research also demonstrates that drugs targeting specific aspects of the pathogenic cascade (e.g., AZM) may have potential clinical utility and elucidates the critical role of glycan and lipid metabolism in CNS development and function.

## Data availability

All data are included in the manuscript and its online supplementary materials. The analytic methods, and study materials will be made available to other researchers for purposes of reproducing the results or replicating the procedure.

## Acknowledgements

The authors thank Dr. Jacques Michaud for stimulating discussions and Dr. Mila Ashmarina for helpful advice and for critical reading of the manuscript. We also thank Steven Lai and Bindesh Shrestha for the essential help and support in the LC-MS/MS and MALDI MSI analyses of mouse brains.

## Supplementary material

Supplementary material is available online.

