## Supplemental for "Mice with mono-allelic p.R37H *Dhdds* variant show aberrant glycosylation and interneuron deficits"

#### Supplementary material

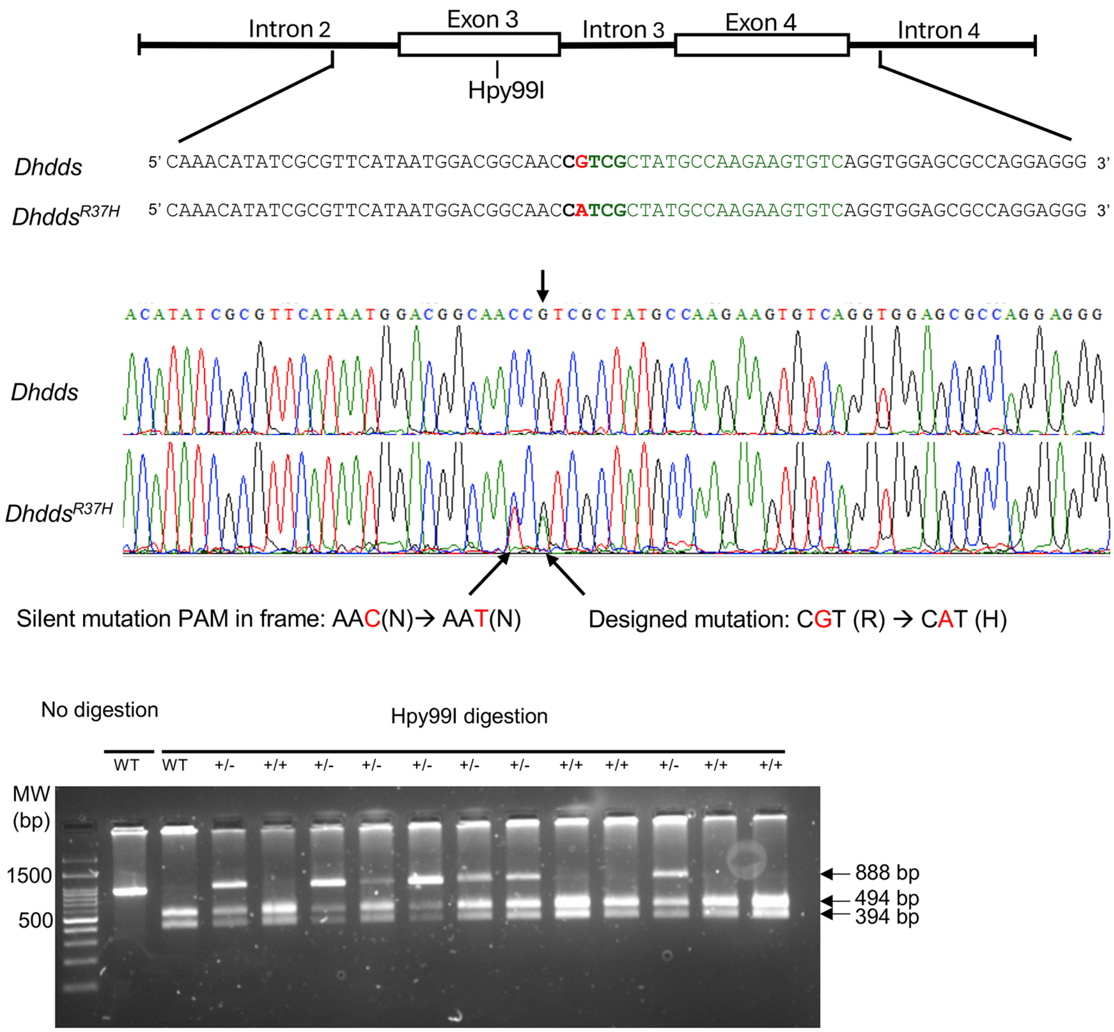


**Supplementary Figure 1. Generation of *Dhdds*$^{R37H}$ mouse strain.**
**(A)** Scheme showing CRISPR-Cas9/sgRNA-targeting site in *Dhdds* exon 3. The sgRNA-targeting sequence is shown in green. The c.50G>A (p.R37H) change is shown in red. The G>A substitution disrupts the Hpy99I restriction site (shown in bold). The exon sequence is capitalized. **(B)** Sanger sequencing of genomic DNA clones showing the genetic variants in exon 3 of the *Dhdds* gene in the *Dhdds*$^{R37H}$ founder mice. The sequences are aligned with the murine *Mus musculus* DNA. **(C)** Genotyping of *Dhdds*$^{R37H+/-}$ mice by PCR/enzymatic restriction assay. The PCR-amplified 888-bp fragment of genomic DNA across the c.50G>A change is cleaved by the Hpy99I restriction endonuclease into 494-bp and 394-bp fragments in the WT but not in *Dhhds*$^{R37H+/-}$ mice showing the undigested 888-bp fragment as well as the 494 and 394 bp fragments.

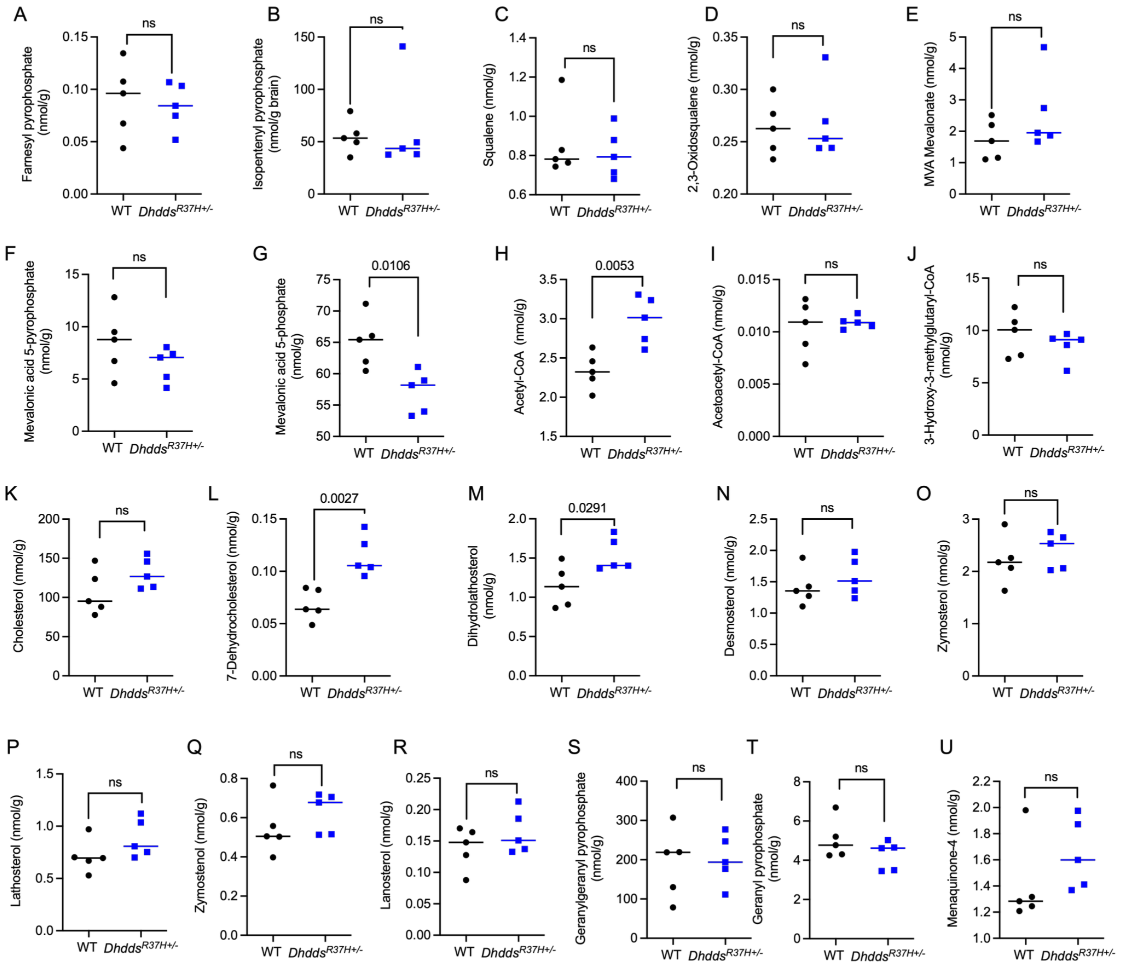


**Supplementary Figure 2. *Dhdds*$^{R37H+/-}$ mice do not present major changes in the brain levels of mevalonate pathway metabolites.**

Levels (nmol/g of wet tissue) of farnesyl pyrophosphate **(A)**, isopentenyl pyrophosphate **(B)**, squalene **(C)**, 2,3-oxidosqualene **(D)**, mevalonate **(E)**, mevalonate 5-pyrophosphate **(F)**, mevalonate 5-phosphate **(G)**, Acetyl-CoA **(H)**, Acetoacetyl-CoA **(I)**, 3-hydroxy-3-methylglutaryl-CoA (HMG-CoA), **(J)**, cholesterol **(K)**, 7-dehydrocholesterol **(L)**, dihydrolathosterol **(M)**, desmosterol **(N)**, zymosterol **(O)**, lathosterol **(P)**, zymostenol, **(Q)**, lanosterol **(R)**, geranylgeranyl pyrophosphate **(S)**, geranyl pyrophosphate **(T)** and menaquinone-4 **(U)** were measured by LC-MRM/MS in water/methanol/chloroform (1:6:2) extracts of brain tissues of *Dhdds*$^{R37H+/-}$ and WT mice ($n$ = 5, 3 males and 2 females for each genotype). All graphs show individual results, means and SD. $P$-values are calculated by two-tailed $t$-test.

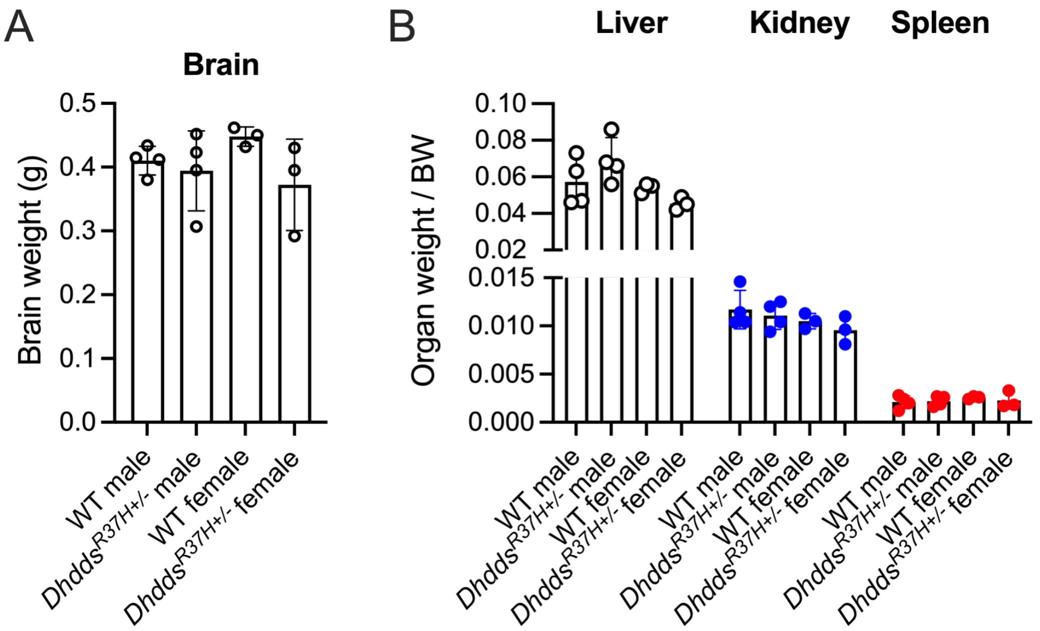


**Supplementary Figure 3. *Dhdds*$^{R37H+/-}$ mice do not present major changes in the brain or organ weigh.**

Wet weight of brains **(A)**, livers, combined kidneys, and spleens **(B)** of 6-month-old *Dhdds*$^{R37H+/-}$ mice and their age and sex matching WT controls was analyzed at sacrifice. Graphs show individual values, means and SD for brains and internal organs, normalised for BW (*n* = 3-4 mice per sex per genotype). Significance of changes was assessed by one-way ANOVA.

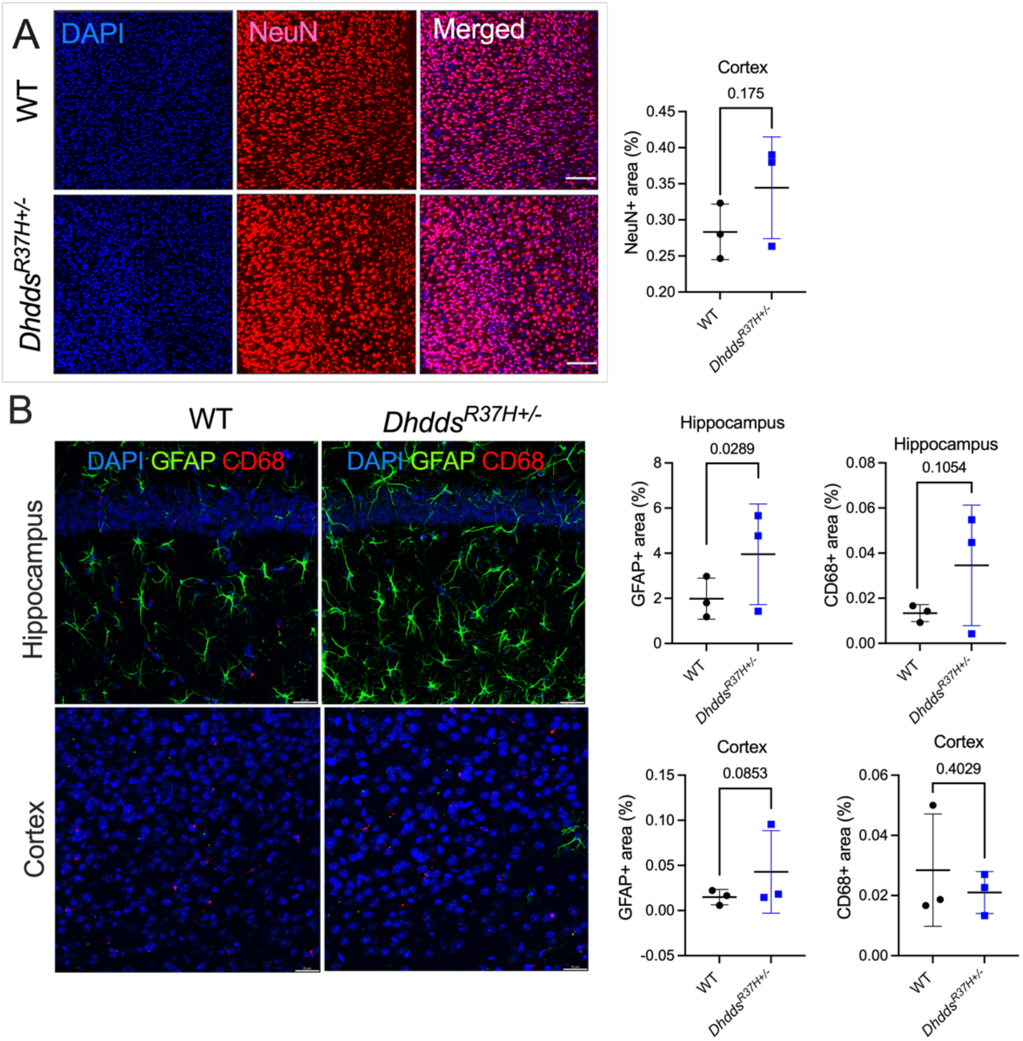


**Supplementary Figure 4. Immunofluorescent microscopy analysis reveals signs of astrogliosis but not of severe neurodegeneration or neuroinflammation in the brains of 8-month-old *Dhdds*$^{R37H+/-}$ mice.**

Panels show representative confocal microscopy images of brain cortices (layers IV-V) and CA1 areas of hippocampus of WT and *Dhdds*$^{R37H+/-}$ mice labeled for **(A)** neurons (NeuN, red), or **(B)** markers of micro- and astrogliosis, CD68+ activated microglia (red) and GFAP+ astrocytes (green). DAPI (blue) was used to stain nuclei. Graphs show quantification of immunofluorescence using ImageJ software. Individual data, means and SD for 3 mice per genotype, 3 areas per mouse, are shown. Statistical analysis was performed by nested two-tailed *t*-test. Size bars are equal to 100 μm **(A)** or 25 μm **(B)**.

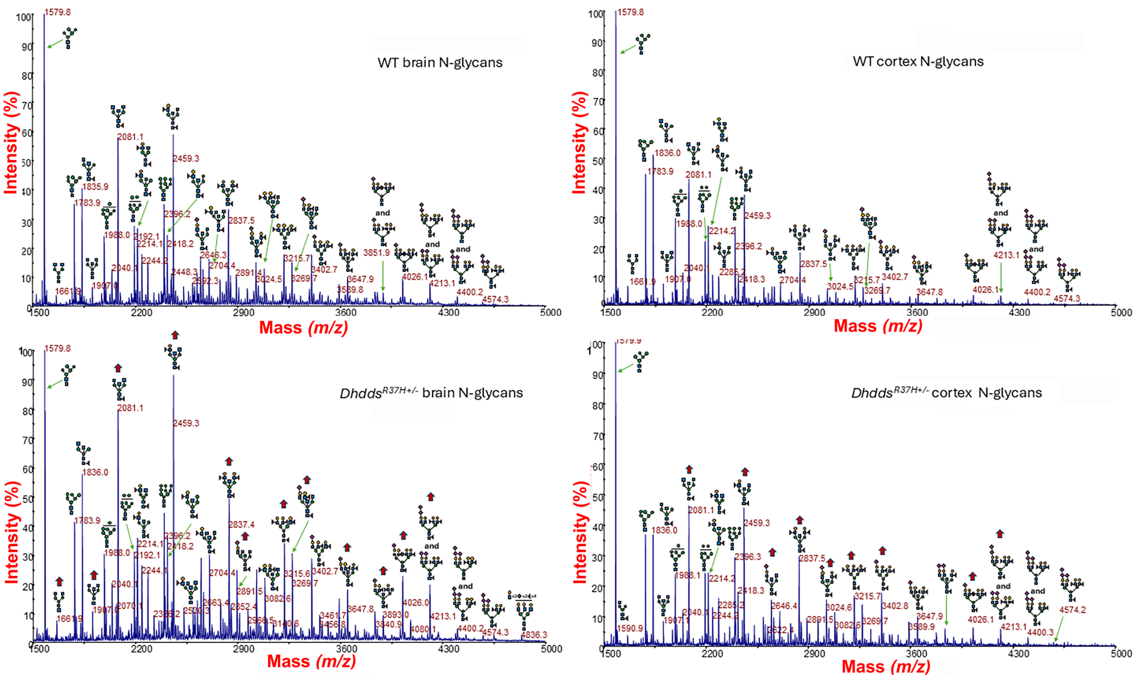


**Supplementary Figure 5. Representative MALDI mass spectra of permethylated N-glycans from brain (A) and cortex (B) of WT and *Dhdds*$^{R37H+/-}$ mice.**

MS profiles are shown in the mass range between *m/z* 1500 and 5000, where glycosylation changes are the most evident. Red arrows mark structures showing major increases in relative peak intensities. GlcNAc, blue square; Man, green circle; Gal, yellow circle; Neu5Ac, purple diamond; Neu5Gc, light blue diamond; Fuc, red triangle.

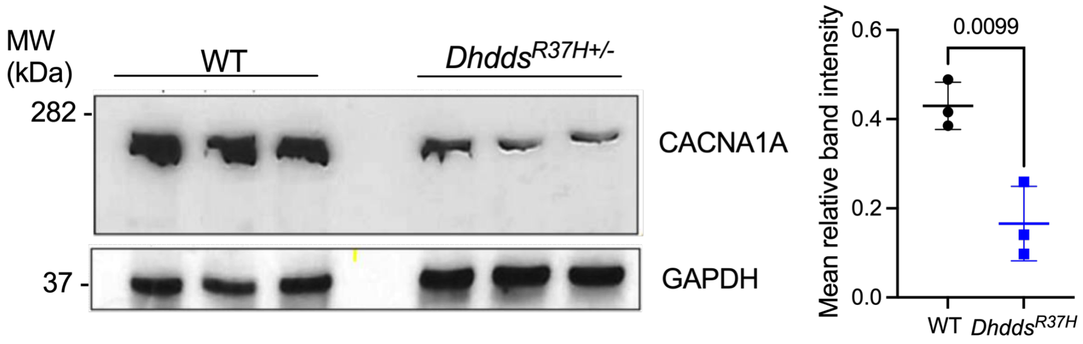


**Supplementary Figure 6. CACNA1A protein levels are reduced in brain tissues homogenates of *Dhdds*$^{R37H+/-}$ compared to WT mice.**

**(A)** Representative immunoblot images and **(B)** quantitative analysis of CACNA1A band intensities normalized by tubulin. Individual results, means and SD from 3 mice per genotype are shown. *P*-values were calculated using two-tailed *t*-test.

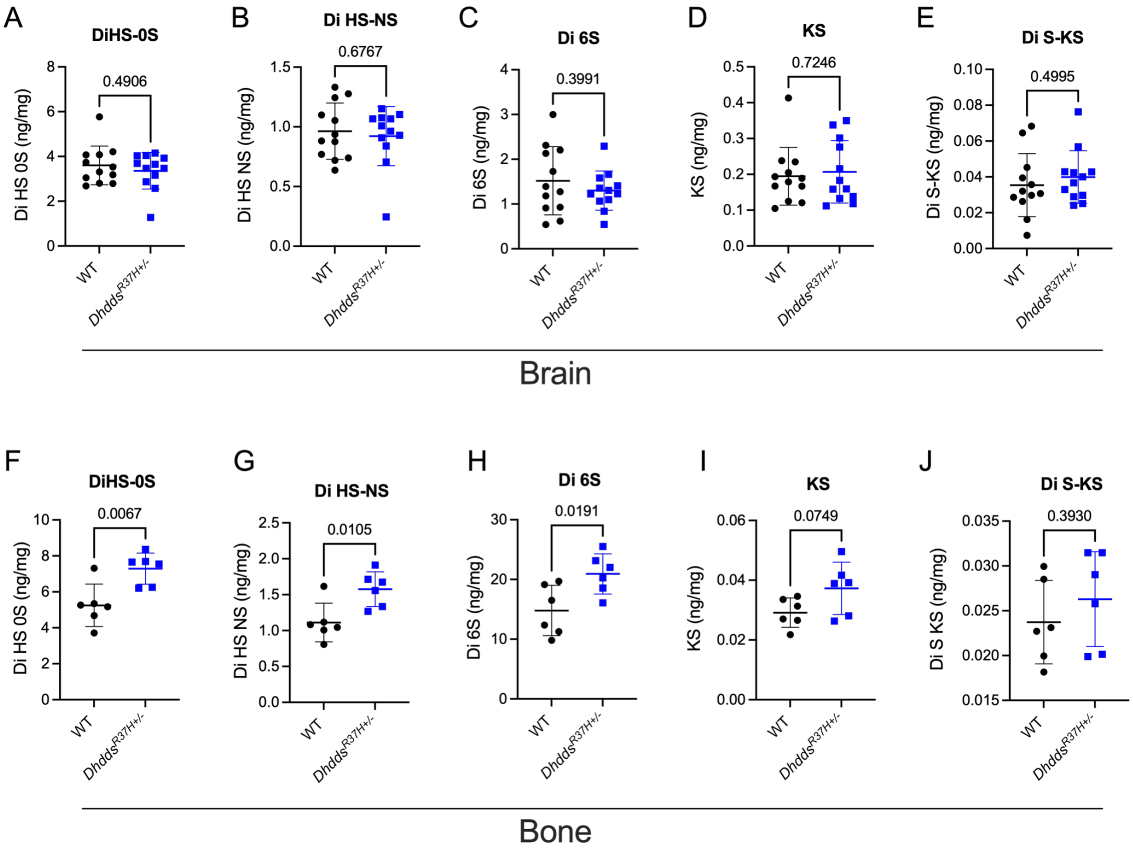


**Supplementary Figure 7. Levels of glycosaminoglycans are increased in the bone tissue of *Dhdds*$^{R37H+/-}$ compared to WT mice.**

Levels of disaccharides produced by enzymatic digestion of heparan sulfate (ΔDiHS-0S and ΔDiHS-NS), chondroitin sulfate (Di 6S) and mono (KS) and di-sulfated keratan sulfate (DiS-KS) were measured by tandem mass spectrometry in brain and tibia bone tissues of WT and *Dhdds*$^{R37H+/-}$ mice at the age of 2, 4 and 6 months. All graphs show individual data, means and SD ($n$ = 6, 3 males and 3 females for each genotype). $P$-values were calculated by two-tailed $t$-test.

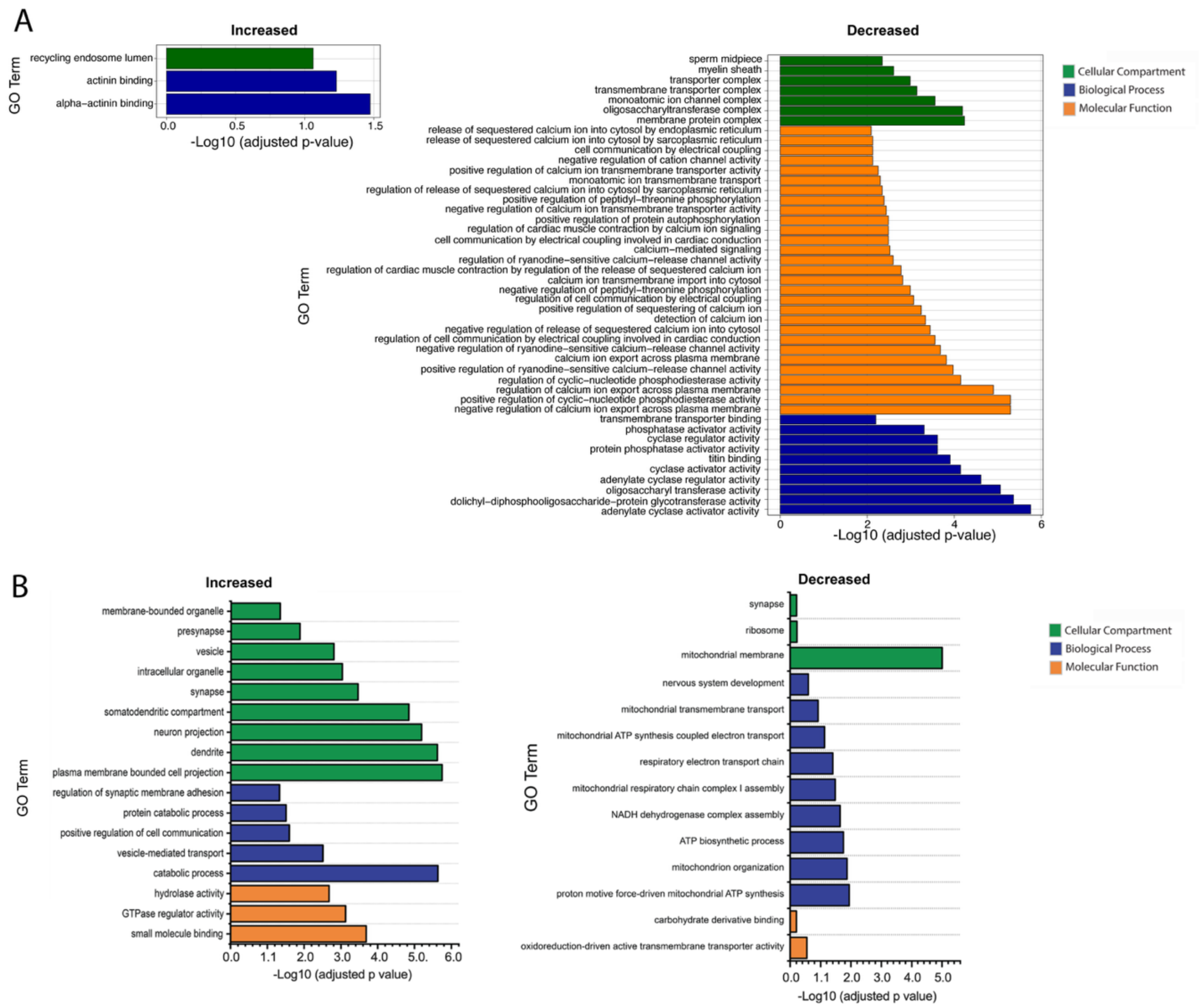


**Supplementary Figure 8. Gene ontology (GO) terms plotted versus significance of change of the proteins altered in the total brains and synaptosomes of *Dhdds*$^{R37H+/-}$ mice compared to WT littermates showing significantly enriched biological processes.**
**(A)** Analysis of total brain proteome showing biological processes downregulated and upregulated in *Dhdds*$^{R37H+/-}$ mice. **(B)** Analysis of purified brain synaptosomes showing biological processes downregulated and upregulated in *Dhdds*$^{R37H+/-}$ mice. The data represent values where the Benjamini corrected *P*-value is below the highest Benjamini corrected *p*-value for the GO terms ($n$ = 5, 3 males and 2 females for each genotype).

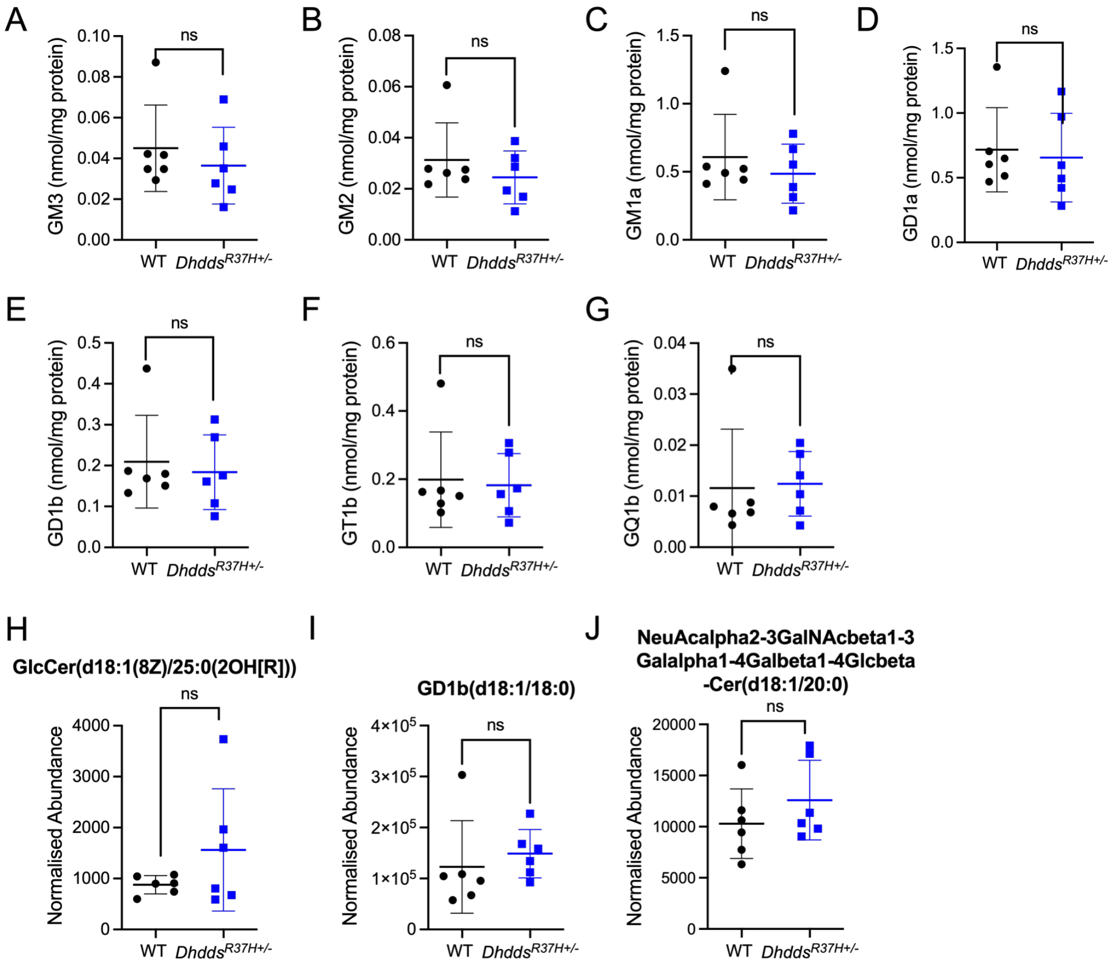


**Supplementary Figure 9.** ***Dhdds*$^{R37H+/-}$ mice do not present major changes in the brain levels of major GSL.**

Levels (nmol/mg of protein) of GM3 **(A)**, GM2 **(B)**, GM1a **(C)**, GD1a **(D)**, GD1b **(E)**, GT1b **(F)** and GQ1b **(G)** gangliosides were measured in total brain extracts by normal HPLC analysis of fluorescently labeled glycans. **(H-J)** Levels of individual GSL species in total brain extracts were estimated by untargeted lipidomic LC-MS/MS analysis ($n$ = 5 mice per genotype). Individual results, means and SD are shown. $P$-values were calculated by two-tailed $t$-test.

**Supplementary table 1A. Detailed list of N-linked structures from whole mouse brain and mouse cortex**

P = Paucimannose; O = Oligomannose; H = Hybrid; C = Complex; B = Bisected; F = Fucosylated; S = Sialylated

| | Theoretical *m/z* | Structure | Glycan type |
|---|---|---|---|
| **1** | 1141.57 | 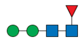 | **P - F** |
| **2** | 1171.58 | 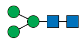 | **P** |
| **3** | 1345.67 | 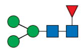 | **P - F** |
| **4** | 1375.68 | 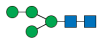 | **P** |
| **5** | 1416.71 | 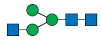 | **C** |
| **6** | 1579.78 | 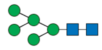 | **O** |
| **7** | 1590.80 | 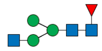 | **C - F** |
| **8** | 1661.83 | 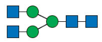 | **C** |
| **9** | 1783.88 | 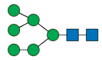 | **O** |
| **10** | 1794.90 | 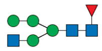 | **H - F** |
| **11** | 1824.91 | 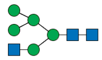 | **H** |
| **12** | 1835.92 | 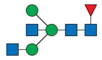 | **C - B - F** |
| **13** | 1865.93 | 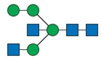 | **H - B** |
| **14** | 1906.96 | 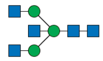 | **C - B** |
| **15** | 1968.99 | 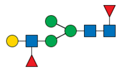 | **C - F** |
| **16** | 1987.98 | 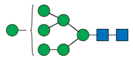 | **O** |
| **17** | 1999.00 | 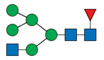 | **H - F** |
| **18** | 2040.02 | 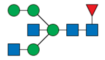 | **H - B - F** |
| **19** | 2040.02 | 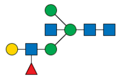 | **C - B - F** |
| **20** | 2040.02 | 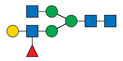 | **C - F** |
| **21** | 2070.03 | 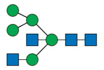 | **H - B** |
| **22** | 2081.05 | 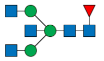 | **C -B - F** |
| **23** | 2111.06 | 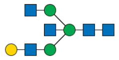 | **C - B** |
| **24** | 2111.06 | 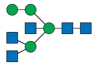 | **H - B** |
| **25** | 2192.08 | 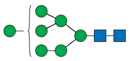 | **O** |
| **26** | 2203.10 | 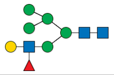 | **H - F** |
| **27** | 2214.11 | 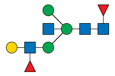 | **C - B - F** |
| **28** | 2214.11 | 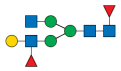 | **C - F** |
| **29** | 2244.12 | 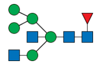 | **H - B - F** |
| **30** | | 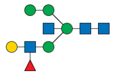 | |

**Supplementary table 1B. N-linked structures found in mouse serum**

| | Theoretical *m/z* | Structure | Glycan type |
|---|---|---|---|
| 1 | 1579.78 | 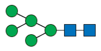 | Oligomannose |
| 2 | 1590.80 | 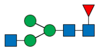 | Truncated (GlcNAc deficient) |
| 3 | 1620.81 | 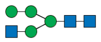 | Hybrid |
| 4 | 1620.81 | 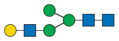 | Truncated (GlcNAc deficient) |
| 5 | 1661.84 | 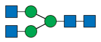 | Truncated (Gal deficient) |
| 6 | 1783.88 | 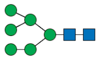 | Oligomannose |
| 7 | 1794.90 | 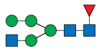 | Hybrid |
| 8 | 1807.89 |  | Truncated (Man deficient) |
| 9 | 1824.91 | 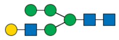 | Hybrid |
| 10 | 1824.91 | 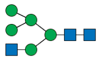 | Hybrid |
| 11 | 1835.93 | 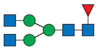 | Truncated (Gal deficient) |
| 12 | 1865.93 | 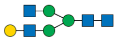 | Truncated (Gal deficient) |
| 13 | 1987.98 | 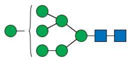 | Oligomannose |
| 14 | 2011.99 | 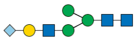 | Truncated (GlcNAc deficient) |
| 15 | 2040.02 | 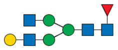 | Truncated (Gal deficient) |
| 16 | 2070.03 | 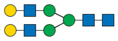 | Truncated (NeuGc deficient) |
| 17 | 2192.08 | 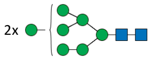 | Oligomannose |
| 18 | 2216.09 | 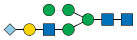 | Hybrid |
| 19 | 2186.08 | 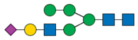 | Hybrid |
| 20 | 2244.12 | 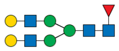 | Truncated (NeuGc deficient) |
| 21 | 2257.12 | 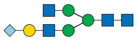 | Truncated (Gal deficient) |
| 22 | 2396.18 | 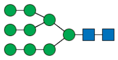 | Oligomannose |
| 23 | 2420.19 | 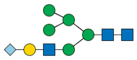 | Hybrid |
| 24 | 2431.21 | 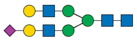 | Truncated (NeuAc deficient) |
| 25 | 2461.21 | 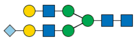 | Truncated (NeuGc deficient) |
| 26 | 2605.30 | 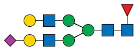 | Truncated (NeuAc deficient) |
| 27 | 2635.31 | 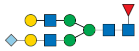 | Truncated (NeuGc deficient) |
| 28 | 2852.40 | 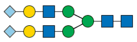 | Sialylated |
| 29 | 3026.49 | 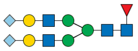 | Sialylated |
| 30 | 3243.59 | 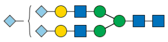 | Sialylated |

### Supplementary methods

#### Animal models

All animal experiments were approved by the Animal care use Committee of Centre Hospitalier Universitaire (CHU) Ste-Justine (approval numbers: 2021-3194 and 2021-7105) and conducted in accordance with the Canadian Council on Animal Care guidelines. Mice were housed in an enriched environment with continuous access to food and water, under constant temperature and humidity, on a 12:12 h light/dark cycle. Mice were kept on a normal chow diet (5% fat, 57% carbohydrate).

The knock-in *Dhdds*$^{R37H}$ C57BL/6 mouse strain, carrying the analog of the human variant c.110G>A:p.R37H in exon 3 of the *Mus musculus* Dehydrodolichyl diphosphate synthase (*Dhdds*) gene, was generated at McGill Integrated Core for Animal Modeling using CRISPR-Cas9 technology. A single-guide nucleotide RNA (sgRNA) was designed, using the CRISPR guide design tools (https://zlab.bio/resources), to target a genomic site on the murine *Dhdds* locus with minimal potential off-target effects. The sgRNA and Cas9 mRNA were microinjected into zygotes together with the single-stranded oligodeoxynucleotide (ssODN), barring a c.50G>A mutation encoding for the R37H change (100 ng/µl of ssODN, 20 ng/µl of sgRNA, and 20 ng/µl of Cas9; Supplementary Fig.1 [40]). After culturing overnight, two-cell zygotes were transferred into surrogate mothers, and pups were delivered at full term. At weaning, pups were genotyped by Sanger sequencing of single-allele fragments obtained by PCR amplification of genomic DNA from the tail clips. The founder mice were bred to C57BL/6n mice to generate F1 heterozygous animals. To test for the presence of potential off-target effects, the sgRNA sequence (GACACTTCTTGGCATAGCGA) was blasted against the *M. musculus* genome. The ~800-bp fragments around top three matches in exon sequences, presenting the highest identity score, were amplified by PCR from genomic DNA of F1 founders followed by Sanger sequencing to analyze for the presence of deleterious variants. The mice were further backcrossed to C57BL/6n mice for three generations to segregate and eliminate potential off-target changes in other alleles. The progenies were intercrossed to generate WT and heterozygous (*Dhdds*$^{R37H+/-}$) animals. Mouse genotyping was performed on tail genomic DNA, extracted using standard protocols. PCR amplifications were performed across the c.50G>A mutation with genomic DNA as template, using the primer set, 5′-CCCTGGGTAGCCAGCAACAA-3′ and 5′-AGGGGAAAGGAGGGGAGGAA-3′ and the following PCR program: 94 °C for 4 min, 35 cycles (94 °C for 30 s, 61°C for 30 s, and 72 °C for 2 min), and 72 °C for 10 min. Since the c.50G>A mutation eliminated the *Hpy99I* restriction site, to distinguish between mutant and WT amplicons, the amplified 888-bp PCR fragment was digested with *Hpy99I* (New England Biolabs, catalog no. R0615S) at 37 °C for 1 h and then separated on a 2% agarose gel. The fragments of 494 and 394 bp were detected for the WT allele, whereas the undigested 888-bp fragment together with the 494- and 394-bp fragments were detected for the heterozygous *Dhdds*$^{R37H+/-}$ mice.

#### Targeted analysis of dolichols and polyprenols in the mouse tissues

Mouse brains were homogenized in deionized water (50 μL per 100 mg of tissue) using a MM 400 mill mixer at 30 Hz for 2 min. Then, the methanol-chloroform mixture (3:1 v/v, 7.5 μL per mg of tissue) containing butylated hydroxytoluene antioxidant (BHT) was added. The samples were homogenized again for 2 min and then sonicated in an ice-water bath for 5 min, followed by centrifugation at 21,000 x *g* for 10 min. 8-μL aliquots of supernatants or serially diluted internal standards (IS) were analyzed by LC-MRM/MS in a positive-ion detection mode using an Acquity UPLC system (Waters) coupled to a QTRAP 6500 mass spectrometer (Sciex). Samples were separated on a C8 column (2.1x50 mm, 2.5 μm) using binary-solvent gradient elution (A phase, 0.1% formic acid in water; B phase, 0.1% formic acid in 1:1 isopropanol-acetonitrile mixture, 80% to 100% B in 15 min) at 0.35 mL/min and 55 °C. Concentrations of the detected compounds were calculated by interpolating the reconstructed peak areas into the calibration curves prepared with standard solutions.

#### Targeted analysis of mevalonate pathway metabolites

For targeted quantitation of the mevalonate (MVA) pathway metabolites by LC-MRM/MS, each mouse brain sample was thawed and homogenized in water-methanol-chloroform mixture (1:6:2 v/v, 10 µL per mg of tissue) using a MM 400 mill mixer for 2 min at 30 Hz, followed by sonication in an ice water bath for 3 min and centrifugation at 21,000 x *g* for 10 min. Internal standards (IS) consisted of serially diluted solutions of d3-labeled mevalonic acid, IPPs, CoAs and HMG-CoA in 50% methanol. 100 µL of the clear supernatant from each sample was dried under a flow of nitrogen, solubilized in 20 µL of IS solution and mixed with 20 µL of 100 mM 3-nitrophenylhydrazine solution and 20 µL of 100 mM EDC/3% pyridine solution. The mixtures were incubated at 50 °C for 30 min. Then, 15 µL aliquots were analysed by UPLC-MRM/MS with a negative-ion detection mode on an Agilent 1290 UHPLC system equipped with a C18 column (2.1x150 mm, 1.8 µm) and coupled to an Agilent 6495B QQQ mass spectrometer. Binary-solvent gradient elution used 0.01% formic acid in water (A) and 0.01% formic acid in acetonitrile (B). Concentrations of detected compounds were calculated by interpolating the reconstructed linear-regression calibration curves with peak areas or peak area ratios recorded with the sample solutions.

#### *Cis*-PTase structural modeling

The molecular dynamics (MD) simulations were performed using the Schrödinger Maestro release 2024-1 (Schrödinger LLC). First, missing loops were added to the structure of h*cis*-PT in complex with FPP (PDB 6Z1N) using MODELLER[41], and the R37H DHDDS mutation was introduced using Schrödinger Maestro release 2024-1 (Schrödinger, LLC). The structure was prepared using the

Protein Preparation Wizard. Missing hydrogen atoms were added reflecting a pH value of 7.2 ± 1.0, followed by optimization of the hydrogen bond network. The system setup tool was used to solvate the systems using the TIP3P model. Potassium or chloride ions were added to neutralize the charge and to obtain a final salt concentration of 150 mM. All MD simulations were performed using Desmond with the OPLS4 force field[42]. The simulations were conducted under a Langevin temperature and pressure control, using periodic boundary conditions with particle-mesh Ewald (PME) electrostatics with a 9 Å cutoff for long-range interactions. The systems were equilibrated using the default relaxation protocol. The production simulations were carried out for 250 ns with a constant pressure of 1 atm and a constant temperature of 300 K starting from a random seed. The results were manually inspected using the Maestro suite. The protein-ligand contacts were analyzed using the simulation interaction diagram tool. Structural illustrations were prepared with PyMol 3.0 (Schrödinger, LLC).

#### Recombinant *cis*-PTase expression and purification

Recombinant DHDDS and NgBR proteins were prepared as previously described[13, 35]. Briefly, *E. coli* T7 express competent cells were co-transformed with $DHDDS^{WT}$ (residues 1-333) or $DHDDS^{R37H}$ and NgBR (residues 73-293), grown in Terrific Broth medium at 37 °C until reaching $OD_{600nm}$ = 0.6 and induced at 16°C by adding 0.5 mM isopropyl β-D-1-thiogalactopyranoside (IPTG). Proteins were expressed at 16 °C for 16-20 hr, harvested by centrifugation (~5,700x*g* for 15 min), and then resuspended in a buffer containing 20 mM 4-(2-hydroxyethyl)-1-piperazineethanesulfonic acid (HEPES), pH 7.5, 150 mM NaCl, 1 mM tris(2-carboxyethyl)phosphine (TCEP) and 0.02% (w/v) triton X-100, supplemented with 1 μg/ml DNase I and a protease inhibitor mixture. Resuspended cells were homogenized and disrupted in a microfluidizer. Soluble proteins were recovered by centrifugation at 40,000x*g* for 45 min at 4 °C. The proteins were purified on a HisTrap HP column, followed by purification on a Strep-Tactin column and TEV protease cleavage of the purification tags and TRX fusions. The reaction mixture was concentrated and loaded onto a Superdex-200 preparative size-exclusion column pre-equilibrated with 20 mM 4-(2-hydroxyethyl)-1-piperazineethanesulfonic acid (HEPES), pH 7.5, 150 mM NaCl, 1 mM tris(2-carboxyethyl)phosphine (TCEP). Purified proteins were flash-frozen in liquid nitrogen and stored at -80 °C until use. Protein purity was >95%, as assessed by SDS-PAGE.

#### *Cis*-PTase enzymatic activity assays

Enzymatic activity of the recombinant *cis*-PTase proteins was measured using a radioligand-based assay as described previously[13, 35, 43]. Briefly, 0.1 μM of purified enzyme (WT or mutant) were mixed with [$^{14}$C]-IPP and FPP to initiate the reaction in buffer composed of 25 mM Tris-HCl, pH 7.5, 150 mM NaCl, 10 mM β-mercaptoethanol, 0.02% v/v Triton-X100, 0.5 mM $MgCl_2$ at 30 °C. 15 mM EDTA (final concentration) was added to quench the reaction and 500 μL of water-saturated 1-butanol was added to extract the reaction products by thorough vortexing. Initial rates were measured by quenching the reaction at <10% substrate consumption. The products, encompassing $^{14}$C, were quantitated using a scintillation counter. The $K_m$ value of FPP or IPP were determined in the presence of 100 μM IPP or 10 μM FPP, respectively. Kinetic constants were obtained by non-linear regression (Michaelis-Menten) equation using Prism 10 (GraphPad Software, LLC).

For the preparation of microsomes, the mouse brains were homogenized using a Potter-Elvehjem homogenizer in 4 volumes of ice-cold 0.1 M Tris-HCl buffer, pH 7.0 containing 0.25 M sucrose and 1 mM EDTA. After centrifugation at 1000×*g* and 4 °C for 10 min, the supernatant was centrifuged again at 12,000×*g* and 4 °C for 20 min. The resulting supernatant was immediately recentrifuged at 100,000×*g* for 45 min at 4 °C. The pellet was washed with and resuspended in the reaction buffer (25 mM Tris-HCl, pH 7.4, 5 mM $MgCl_2$, 1.25 mM DTT and 2.5 mM sodium orthovanadate) and homogenized by sonication followed by protein concentration assay. The enzyme assay mixture containing 250 μg of protein in the 1000 μl of the reaction buffer supplemented with 25 μM FPP, 50 μM [$^{14}$C]-IPP (55 mCi/mmol), 10 μM Zaragozic acid A and 0.35% Triton X-100 was incubated at 37 °C for 2 h. The polyprenol pyrophosphate products were extracted with chloroform/methanol (3:2) mixture, followed by washing the organic phase three times with 10 mM EDTA in 0.9% NaCl. The concentration of [$^{14}$C]-labeled products in the organic phase was measured by a scintillation counter.

#### Video-Electroencephalographic recordings

The presence of epileptiform activity was assessed by intracranial video-EEG recordings as described[44, 45]. Intracranial electrode implantation surgeries were performed at P60-P75 under isoflurane anesthesia (5% induction, 2% maintenance adjusted as needed, oxygen flow rate 1 L/min). Buprenorphine (0.10 mg/kg) was administered subcutaneously for post-implantation analgesia. After shaving and cleaning the scalp at the surgical site with povidone-iodine and 70% alcohol, a sagittal anteroposterior scalp incision was made to expose the cranial surface. PFA coated tungsten recording electrodes (A-M Systems, 50.8 μm diameter, Cat 795500), adapted to a custom fabricated 18-channel Mill-Max pedestal were inserted bilaterally in somatosensory cortex (AP/ML/DV: - 1.0/±1.5/-1 mm) and hippocampus CA1 (-1.5/±1.5/2.0 mm) (coordinates relative to Bregma at brain surface, 2 electrodes at each recording site with 2 EMG electrodes in the neck muscle). Reference and ground electrodes were placed over the cerebellum. The electrode constructs were fixed to the skull using optibond (Kerr Dental) and dental cement (Lang Dental). Video-EEG recordings were conducted continuously for 14 days in mutant mice and 10 days in WTs, starting a minimum of 48 h after surgery. Subjects were connected to a headstage preamplifier and to the subsequent Digital Lynx SX (NeuraLynx) or RHD2000 (Intan Technologies) systems, with a sampling rate of 4kHz, and filtered at HP 0.1Hz and LP 1000 Hz on both systems.

All EEG signals were manually verified. Seizures (more than 4 sec associated with or without behavioral component) were identified and reviewed by 3 experimenters. Behavioral states (non-rapid eye movement sleep, rapid eye movement sleep and wakefulness) were identified manually based on the EEG, spectral power, and EMG. Power spectral density color map was calculated using the Welch's power spectral density estimate with 5 sec window size and overlapping window of 3 sec.

#### Pentylenetetrazol-induced seizure threshold and acetazolamide administration

Following baseline EEG recordings, seizure susceptibility to pentylenetetrazol (PTZ) was evaluated with a continuous intravenous infusion (dissolved in normal saline; 10 mg/ml) injected in the tail vein at 0.1 ml/min as previously described[45]. PTZ infusions were stopped with the appearance of the first seizure activity on EEG. The seizure threshold for each animal was defined as the minimal PTZ dose (mg/kg BW) required for seizure induction.

In a separate set of animals, pre-treatment with acetazolamide (AZM) was conducted before PTZ infusion. A stock solution of AZM (Sigma-Aldrich, #A6011) was prepared by dissolving the drug powder in DMSO at a concentration of 200 mg/mL. This was then diluted to an AZM concentration of 2 mg/mL using saline. The experimental group of *Dhdds*$^{R37H+/-}$ mice received an AZM injection (40 mg/kg) [47, 48]. The control *Dhdds*$^{R37H+/-}$ and littermate WT mice were injected with saline. Injections were administered intraperitoneally (i.p.) 45 min prior to the infusion of PTZ. PTZ seizure threshold was analyzed as described above.

#### Patch Clamp

The functional impact of the *Dhdds*$^{R37H+/-}$ variant was assessed using voltage-clamp recordings of spontaneous excitatory/inhibitory postsynaptic currents (sEPSCs/sIPSCs) and miniature (mIPSC) inhibitory postsynaptic currents in layer V principal cells from the primary somatosensory cortex (S1) of *Dhdds*$^{R37H+/-}$ mice and littermate controls.

**Slice preparation**. P26-P35 mice of both sexes were deeply anesthetized with isoflurane and intracardially perfused with ~20 mL of carbogenated (5% $CO_2$ balanced with $O_2$) choline chloride cutting solution containing (in mM): 132.5 choline chloride, 2.5 KCl, 0.7 $CaCl_2$, 25 $NaHCO_3$, 8 glucose, 3 $MgCl_2$, 1.2 $NaH_2PO_4$, pH 7.2-7.4, 330–340 mOsm/L, ~4°C. The brain was then extracted and acute 300 µm-thick coronal slices were made with a vibratome (Leica VT1000S) in carbogenated (5% $CO_2$ balanced with $O_2$) choline chloride cutting solution (~4°C). The slices were maintained at room temperature in artificial cerebrospinal spinal fluid (ACSF) composed of (in mM): 125 NaCl, 2.5 KCl, 1.25 $NaH_2PO_4$, 26 $NaHCO_3$, 2 $MgSO_4$, 2 $CaCl_2$, and 10 D-glucose; 305-315 mOsm/L, pH 7.2-7.4, and continuously infused with carbogen (5% $CO_2$ balanced with $O_2$). During experiments, slices were continuously perfused (2 mL/min) with the same ACSF (31.5°C) in the recording chamber.

**Voltage-clamp.** Glass recording electrodes (BF150-86-10, Sutter Instrument, Novato, CA, US) were used with a resistance ranging from 3.0–5.0 MΩ when filled with cesium-based intracellular solution of the following composition (in mM): 130 $CsMeSO_4$, 5 CsCl, 2 $MgCl_2$, 10 phosphocreatine, 10 HEPES, 0.5 EGTA, 4 ATP-Tris, 0.4 GTP-Tris, 0.5-1% biocytin, 2 QX-314, pH 7.2-7.3 (adjusted with CsOH), 280–295 mOsm/L. Somatic whole-cell voltage-clamp recordings of S1 cortical layer V principal neurons were performed. Recordings of sEPSC were performed in voltage-clamp mode holding at –70 mV. Recordings of sIPSC and mIPSCs were performed in voltage-clamp mode holding at +10 mV in the presence of NBQX (10 µM; Abcam Biochemicals) and DL-AP5 (100µM; Abcam Biochemicals); with the addition of TTX (1 µM) (Alomone Labs) for mIPSCs. mIPSC recordings were obtained after minimum 7 min of perfusion with solution containing NBQX, DL-AP5, and TTX. Series resistance in voltage-clamp was monitored throughout the experiment and cells that had substantial increases in series resistance (>15%), or an access resistance above 18 MΩ, during recording were discarded. Data acquisition was performed using the Multiclamp 700B amplifier (Molecular Devices, CA, US), low-pass filtered at 2 kHz, digitized (sampling rate 10kHz; Digidata 1440, Molecular Devices) and recorded by a computer with Clampex 11.4 software (Molecular Devices).

**Electrophysiological data analysis.** Analysis of electrophysiological recordings was performed using MATLAB (R2025a). For the analysis of sEPSCs, sIPSCs and mIPSCs, 2.5 minutes were recorded at each condition. Traces were first low-pass filtered at 40 Hz. A 10s trace with the lowest standard deviation was picked as the baseline. Events with a minimum prominence of 2.5 standard deviations away from the mean of the baseline period were detected throughout the traces as events and then manually verified by at least 2 experimenters. The amplitude, half-amplitude duration and inter-event interval were calculated on the filtered trace. The remaining analysis was done using the original traces with event time tags generated from the filtered traces. The 20–80% rise time of the events was determined from a second degree polynomial fit of the rising phase. The decay time constant of the detected events were determined from the exponential fit (100–37%) of the decaying phase. Charge transfer was calculated by integrating the area under each EPSC or IPSC event and then averaged over all events. The ΔQ *f was calculated as the charge transfer of the averaged PSC (ΔQ) multiplied by the PSC rate. Graphpad Prism was used to generate figures, and Welch's t-test was used for statistical analysis.

**Immunohistochemistry and anatomical identification.** For post hoc anatomical identification, each recorded neuron was filled with biocytin (0.5-1%, Sigma) during whole-cell recordings (minimum 15 min). After fixing in 4% PFA for a minimum of 2 days, slices were permeabilized with 0.3% Triton X-100 and incubated at 4°C with streptavidin-conjugated Alexa-488 (1:1000, Invitrogen, Cat# S11223) in TBS to visualize biocytin-filled cells. Slices were then incubated with DAPI for 10 min before mounting. Cells were imaged using a Leica SP8 confocal microscope (PIM facility, CHUSJ). A z-stack image was taken for each cell under 20x magnification with a z-step size of 1 µm. Images were evaluated using the Leica LAS X program. Principal cell identity was confirmed by a combination of cell morphology and biophysical properties.

#### Untargeted lipidomic analysis

Mouse brain hemispheres were homogenised using Kinematica polytron PT 3000 homogenizer in methanol/water mixture (1:1 v/v, 1.5 ml per 50 mg of tissue) followed by centrifugation at 16,000x*g* for 10 min. The solid precipitate was homogenised in dichloromethane/methanol 3:1 mixture followed by centrifugation, and the supernatant (organic extract) dried under the flow of nitrogen. The organic extract was analyzed by LC-MS/MS on UPLC-MRT system (Waters) using CSH C18 Acquity Premier column (2.1×100 mm, 1.7 μm) eluted at 0.4 ml/min and 55°C. The mobile phase A was 600/390/10 (Acetonitrile/Water/1 M aqueous ammonium formate), and B was 900/90/10 (IPA/ACN/1 M aqueous ammonium formate). The gradient program was as follows: initial 50% B; 0-0.5 min 50-53% B; 0.5-4 min 53-55% B; 4-7 min 55-65% B, 7-7.5 min 65-80% B; 7.5-11 min 80-99% B; 11-12 min 99-50% B. The acquisition was conducted in the positive mode. Identification of metabolites and quantification was done using Progenesis QI software by integration of areas under chromatograms. All data were normalized to those obtained with pulled extracts.

### Quantitative analysis of brain glycosphingolipids

GSLs were analyzed essentially as described previously[49]. Lipids from aqueous mouse brain homogenates (~2.5 mg in 0.5 ml) were extracted with chloroform: methanol (1:2, v/v) overnight at 4 °C. The GSLs were further purified using solid-phase C18 columns (Telos; Kinesis). After elution, the GSL fractions were dried down under a stream of nitrogen at 42 °C and digested with recombinant endoglycoceramidase (rEGCase I, prepared by Genscript) to release the oligosaccharide headgroups. The liberated free glycans were fluorescently labeled with anthranilic acid (2AA). To remove excess 2AA label, labeled glycans were purified using DPA-6S SPE columns (Supelco). Purified 2AA-labeled oligosaccharides were separated and quantified by NP-HPLC as previously described [50]. The NP-HPLC system consisted of a Waters Alliance 2695 separations module and an in-line Waters 2475 multi λ-fluorescence detector set at excitation λ 360 nm and emission λ 425 nm. As a solid phase, a 4.6 × 250-mm TSK gel-Amide 80 column (Tosoh Bioscience) was used. A 2AA-labeled glucose homopolymer ladder (Ludger) was included to determine the glucose unit (GU) values for the HPLC peaks. Individual GSL species were identified by their GU values and quantified by comparison of integrated peak areas with a known amount of 2AA-labeled BioQuant chitotriose standard (Ludger). Protein concentration in homogenates was determined using the BCA assay (Sigma-Aldrich).

#### MALDI MS imaging

Brains of WT and *Dhdds*$^{R37H+/-}$ mice were embedded in 3.6% carboxymethylcellulose (CMC), frozen, and stored at -80 °C. The frozen tissues were cut into 10 μm-thick sagittal sections using a Thermo Scientific cryostat at -10 °C. The sections from WT and *Dhdds*$^{R37H+/-}$ mice of the same sex were placed on the same slide. Frozen tissue sections were places on the glass slides and dried in a desiccator for 15 min before deposition of 5-chloro-2-mercaptobenzothiazole (CMBT) matrix using an HTX M5 sprayer. CMBT was dissolved at a concentration of 15 mg/mL in 90% acetone:10% $ddH_2O$ mixture and sonicated for 10 min. M5 HTX sprayer conditions were as follows: 1100 mm/min nozzle velocity, 1.5 mm track spacing, 0.1 mL/min flow rate, CC pattern, 4 passes, 40 mm nozzle height, 10 psi for nitrogen flow, nozzle temperature to 30 °C, and heated tray temperature 35 °C. MALDI imaging was performed on a MRT mass spectrometer connected to a MALDI source (Waters). Mass spectra were collected by running a series of scans across the tissue sections and integrated using Mass Lynx with *m/z* 100-2.400 mass range in negative ion mode and at 35 μm pixel resolution. The phosphatidylinositol (PI) was used as the mass-to-mass lock correction using *m/z* 885.5499 (-) as reference. Before viewing results, the data sets were subjected to continuous lock mass correction (CLMC) by phosphatidylcholine. Next, high-definition imaging (HDI) software was used to process and convert the MS data into ion imaging format. Signal intensities were normalized to the total ion current in the HDI software. Lipid annotation was also manually curated using HMDB, Lipid Maps and literature reporting MALDI-MS imaging of brain tissue. The 1,000 most intense peaks were acquired in HDI software to visualize at MS resolution of 20,000.

#### Analysis of N-linked glycans

Mouse brain tissue was homogenized and lysed in a chloroform/ methanol/water 4:8:3 (v/v/v), as described previously[51]. The protein pellet was separated from the lipid-containing supernatant by centrifugation and cleaned by repeated washes with acetone/ water (4:1) at 4 °C. The final pellet was dried under a stream of nitrogen and stored at −20 °C until the analysis. The mouse serum was prepared from the blood obtained by cardiac puncture and lyophilized. For *N*-glycan analysis, ~2 mg of each protein sample was resuspended in 0.2 ml of 0.1% RapiGest Surfactant (Waters Corporation, Milford, MA) in 50 mM $NH_4HCO_3$ using an ultrasonic processor equipped with a 2-mm probe (130 W, 50% amplitude, 5 min in pulsing mode). The homogenate was incubated at 100 °C for 5 min, followed by reduction of proteins by 5 mM dithiothreitol (Sigma- Aldrich) at 56 °C for 30 min and alkylation in 15 mM iodoacetamide (Sigma-Aldrich) in the dark at room temperature for 45 min. Glycan chains were cleaved by peptide-*N*-glycosidase F (4 U; Roche Molecular Biochemicals, Mannheim, Germany) overnight at 37 °C. The released *N*-glycans were purified and permethylated by $ICH_3$ in a dimethyl sulfoxide/NaOH slurry, as described[52]. MALDI-TOF and MALDI-TOF/TOF analyses of permethylated *N*-glycans were performed using 5-chloro-2-mercaptobenzothiazole (10 mg/ml in 80:20 methanol/water, v/v) as matrix and acquired on a 4800 proteomic analyzer (AB Sciex) in positive polarity and in reflector mode and detected as sodiated $[M+Na]^+$ions [52]. Data were analyzed using DataExplorer 4.9 software. Glycan structures were assigned based on molecular weight, biosynthetic pathway, and MS/MS spectra using the bioinformatic tools developed by the Consortium for Functional Glycomics (http://functionalglycomics. org).

#### Sample preparation for proteomics and glycoproteomics

Brain samples were lysed in 5% SDS (in 100 mM triethylammonium bicarbonate buffer) using a Bioruptor sonication device (Hologic). Protein concentrations in the lysates were measured using the Pierce BCA Protein Assay Kit (Thermo Fisher Scientific). Sample aliquots with equal amounts of protein were first reduced using 20 mM dithiothreitol (DTT) for 10 minutes at 95 °C and then alkylated with 100 mM iodoacetamide (IAA) for 30 minutes in the dark at room temperature. Samples were subsequently digested with trypsin (Worthington) using S-Trap midi cartridges (ProtiFi) per the manufacturer's instructions. The resulting peptides were labeled with tandem mass tags (TMT) (Thermo Fisher Scientific) and subsequently pooled for a multiplex MS analysis as per the manufacturer's protocol.

#### Fractionation and glycopeptide enrichment

The pooled TMT-labeled peptides were split into two aliquots. The first aliquot (~20% of the total) was resuspended in solvent A (5 mM ammonium formate, pH 9) and fractionated by basic pH reversed phase liquid chromatography (bRPLC) on a reversed phase Waters C18 column (5 µm, 4.6 × 100 mm column) using an increasing gradient of solvent B (5 mM ammonium formate, pH 9, in 90% acetonitrile) on the Ultimate 3000 UHPLC system. Ninety-six fractions were collected and subsequently concatenated into 12 fractions for the proteomics experiment. The second aliquot (~80%) was used to enrich glycopeptides by size exclusion chromatography [53]. Peptides were resuspended in 100 µL of 0.1% formic acid and injected into Superdex peptide 10/300. The peptides were separated using an isocratic flow of 0.1% formic acid for 130 min and early fractions were collected starting at 10 min after injection. These fractions were subsequently concatenated into 12 fractions for the glycoproteomics experiment.

#### Proteome and glycoproteome analysis by liquid chromatography-tandem mass spectrometry (LC-MS/MS)

LC-MS/MS analysis of fractionated samples from both proteomics and glycoproteomics was carried out as described previously with slight modifications[54, 55]. The samples were analyzed on an Orbitrap Eclipse mass spectrometer equipped with Ultimate 3000 liquid chromatography system (Thermo Fisher Scientific Inc.). The peptides/glycopeptides were separated on an analytical column (EasySpray 75 µm × 50 cm, C18 2 µm, 100 Å, with a flow rate of 300 nL/min with a linear gradient of solvent B (100% ACN, 0.1% formic acid) over a 155 min gradient. Precursor ions were acquired at a resolution of 120,000 (at m/z 200), and fragment ions, at a resolution of 30,000 (at m/z 200). Precursor ions were acquired in the Orbitrap mass analyzer in m/z range of 350-1,700 for proteomics and 375-2,000 for glycoproteomics. The fragmentation was carried out using higher-energy collisional dissociation (HCD) method with normalized collision energy of 35 for proteomics or stepped HCD (15, 25, 40) for glycoproteomics. The scans were acquired in top-speed method with 3 sec cycle time between MS and MS/MS.

#### Proteomics and glycoproteomics data analysis

The peptides were identified using Sequest search engine in Proteome Discoverer 3.0 against the Uniprot Human or Mouse protein sequences. The glycopeptides were analyzed using the publicly available software pGlyco3, which uses a built-in human/mouse N-glycan database[56]. Two missed cleavages were allowed for both proteomics and glycoproteomics analysis. For proteomics data, error tolerance for precursor and fragment ions was set to 10 and 0.02 Da ppm, respectively, and for glycoproteomics data it was set to 10 ppm and 20 ppm, respectively. Cysteine carbamidomethylation, TMT mass on lysines and peptide N-termini were set as fixed modification with oxidation of methionine as a variable modification. False discovery rate (FDR) was set to 1% at the peptide-spectrum matches (PSMs), peptide, protein, and glycopeptides levels. For proteomics, quantitation of peptides across different groups was performed using TMT reporter ion intensities with the "reporter ion quantifier" node. To quantify glycopeptides, reporter ion quantification was performed for glycoproteomics raw files in Proteome Discoverer version 3.0 and glycopeptide IDs obtained from pGlyco3 were matched with quantitation on a scan-to-scan basis (MS/MS). A two-sample Student's *t*-test with unequal group variance was used to identify differentially expressed proteins and glycopeptides in the *Dhdds*$^{R37H+/-}$ mice. Statistical analysis was performed using the publicly available computational platforms, Perseus[57], MetaboAnalyst[58], and R studio. Gene Ontology analysis of proteins corresponded to differentially expressed glycopeptides to identify functionally relevant biological processes dysregulated in the *Dhdds*$^{R37H+/-}$ mice.

#### Proteomic analysis of mouse brain synaptosomes

Synaptosomes were isolated from the freshly collected brains of 4-month-old *Dhdds*$^{R37H+/-}$ and WT mice using a commercially available kit (Thermo scientific # 87793) as recommended by the manufacturer. Semi-quantitative label-free proteomic analysis was performed using liquid chromatography–tandem mass spectrometry (LC-MS/MS). Samples were reconstituted in 50 mM ammonium bicarbonate/8 M urea, vortexed and further diluted 1:8 with 10 mM TCEP[Tris(2-carboxyethyl)phosphine hydrochloride; Thermo Fisher Scientific], and vortexed for 1 h at 37 °C. Chloroacetamide (Sigma-Aldrich) was added for alkylation to a final concentration of 55 mM. Samples were vortexed for another hour at 37 °C. One microgram of trypsin was added, and digestion was performed for 8 h at 37 °C. Samples were dried down and solubilized in 5% ACN-4% formic acid (FA). The samples were loaded on a 1.5 µl pre-column (Optimize Technologies, Oregon City, OR). Peptides were separated on a home-made reversed-phase column (150-µm i.d. by 200 mm) with a 106-min gradient from 10 to 30% ACN-0.2% FA and a 600-nl/min flow rate on an Easy nLC-1200 connected to a Exploris 480 (Thermo Fisher Scientific, San Jose, CA). Each full MS spectrum acquired at a resolution of 120,000 was followed by tandem-MS (MS-MS) spectra acquisition on the most abundant multiply charged precursor ions for 3s. Tandem-MS experiments were performed using higher energy collision dissociation (HCD) at a collision energy of 34%. The data were processed using PEAKS 11 (Bioinformatics Solutions, Waterloo, ON) and a Uniprot Mouse database. Mass tolerances on precursor and fragment ions were 10 ppm and 0.01 Da, respectively. Fixed modification was carbamidomethyl (C). Variable selected posttranslational modifications were acetylation (N-ter), oxidation (M), deamidation (NQ), phosphorylation (STY). Protein false discovery rate was set to 1% or less (FDR ≤ 1%).

#### Omics data analysis.

Proteomic and glycoproteomic data were analyzed using R (Version 4.4.1) and R studio (2024-04-2). The following R packages were used for analysis: dplyr (1.1.3), ggplot2 (3.4.3), org.mm.Eg.db (3.19.1). To filter for synaptic proteins in Figure 3D, genes associated with the GO term 'synapse' (GO :0045202) were extracted and matched with the glycoproteomics dataset. All synaptic glycoproteins which were significantly different between *Dhdds*$^{R37H+/-}$ and WT mice (*p*-value <0.001) were visualized in Figure 3D. In Figure 3E, proteins of interest were highlighted, if they were significantly different for *Dhdds*$^{R37H+/-}$ and WT mice (*P*-value <0.05 for oligomannose glycans and *p*-value <0.001 for complex/hybrid glycans). To filter for lipases and lipid esterases, all genes associated with the GO term 'lipase activity' (GO:0016298) were extracted.

Statistical overrepresentation analysis was performed using g:profiler [59]. Differentially expressed genes were selected based on *p*-values <0.01 for proteomics analysis. The GO Ontology was used to identify significantly overrepresented pathways.

#### Behavioral tests

The spontaneous alternation behavior, spatial working memory, and exploratory activity of mice were evaluated using a white Y-maze test as previously described[40]. The maze consisted of three identical white Plexiglas arms (40 × 10 × 20 cm, 120° apart) under dim lighting conditions. Each mouse was placed at the end of one arm, facing the center, and allowed to explore the maze for 8 min. All experiments were performed at the same time of day and by the same investigator to avoid circadian and handling bias. Sessions were video-recorded, and arm entries were scored by a trained observer unaware of the mouse genotype or treatment. Successful alternation was defined as consecutive entries into a new arm before returning to the two previously visited arms. Alternation was calculated as (number of alternations/total number of arm entries − 2) × 100.

NOR test was used for assessing short-term recognition memory[60]. Mice were placed individually in a 44 × 33 × 20-cm (length × width × height) testing chamber with white Plexiglas walls for a 10-min habituation period and returned to their home cage. The next day, mice were placed in the testing chamber for 10 min with two identical objects (red plastic towers, 3 × 1.5 × 4.5 cm) and returned to the home cages, and 1 h later, were placed back into the testing chamber in the presence of one of the original objects and one novel object (a blue plastic base, 4.5 × 4.5 × 2 cm) for 10 min. After each mouse, the test arena as well as the plastic objects were cleaned with 70% ethanol to avoid olfactory cue bias. The discrimination index (DI) was calculated as the difference in the exploration time between the novel and old object divided by total exploration time. A preference for the novel object was defined as DI significantly >0 [61]. Mice who showed a side preference, noted as DI ± 0.20 during the familiarization period, and those who had a total exploration time <3 s were excluded from analysis. For the rotarod test, mice were placed on an automated accelerated Rotarod (Ugo Basile) with a speed increasing from 10 to 50 rpm over 5 min. The time to fall was recorded in 3 test trials performed on 3 consecutive days. A separate group of *Dhdds*$^{R37H+/-}$ mice and their WT siblings received i.p. injections of saline or AZM (40 mg/kg BW, in saline) 45 mins before each rotarod test.

AZM solution was prepared and administered as described above for the PTZ-induced seizure experiment.

#### RNA extraction and sequencing

Total RNA was extracted from ~30 mg of hippocampal tissue using the RNeasy Mini Kit (Qiagen), according to the manufacturer's instructions. RNA was quantified using the NanoDrop 8000. Samples with a 28S/18S ratio >1.8, an OD 260/280 ratio >1.9 and an RNA integrity number >9 (Agilent Bioanalyzer 2100) were chosen for cDNA library construction. Library preparation and sequencing were performed at the Genomics Platform of the Institute for Research in Immunology and Cancer, Montreal. All cDNA libraries were sequenced using single-end strategy (40 M reads per sample) on an Illumina NextSeq500 platform with a read length of 75 bp (single-end mode). Raw data were converted to FASTQ files using bcl2fastq (v2.20) and allowed 0 mismatches in the multiplexing barcode. Data were analyzed by BioJupies using default parameters. Gene expression–level differences were accepted as statistically significant if they had a $p$ value <0.05, and the result was visualized by an in-house Matlab code. The data are available on: https://amp.pharm.mssm.edu/biojupies/notebook/y4cudhb0o.

#### Analysis of glycosaminoglycans by LC-MS/MS

Analysis of glycans in the bone and brain tissues was conducted as previously described[62]. Briefly, 30–50 mg of tissue was homogenized in ice-cold acetone and centrifuged at 12,000×*g* for 30 min at 4 °C. The pellets were dried, resuspended in 0.5 N NaOH and incubated for 2 h at 50 °C. Then, the pH of the samples was neutralized with 1 N HCl, and NaCl was added to the reaction mix at a final concentration of 3 M. After centrifugation at 10,000×*g* for 5 min at room temperature, the supernatants were collected and acidified using 1 N HCl. Following another centrifugation at

10,000×*g* for 5 min at room temperature, the supernatants were collected and neutralized with 1 N NaOH to a pH of 7.0. The samples were diluted at a ratio of 1:2 with 1.3% potassium acetate in absolute ethanol and centrifuged at 12,000×*g* and 4 °C for 30 min. The pellets were washed with ice-cold 80% ethanol, dried at room temperature, and dissolved in 50 mM Tris-HCl buffer. The samples were placed in AcroPrepTM Advance 96-Well Filter Plates with Ultrafiltration Omega 10 K membrane filters (PALL Corporation, New York, NY, USA) and digested with chondroitinase ABC, chondroitinase B, heparitinase, and keratanase II, overnight at 37 °C. The samples were analyzed by mass spectrometry using a 6460 Triple Quad instrument (Agilent technologies, Santa Clara, CA, USA) with Hypercarb columns, as described[62].

### Western blotting

The cerebral cortical tissues were homogenized in five volumes of RIPA lysis buffer (50 mM Tris-HCl, pH 7.4, 150 mM NaCl, 1% NP-40, 0.25% sodium deoxycholate, 0.1% SDS, 2 mM EDTA, 1 mM PMSF), containing protease and phosphate inhibitor cocktails (Sigma # 4693132001 and # 4906837001), using a Dounce homogenizer. The homogenates were kept on ice for 30 min and centrifuged at 13,000x*g* and 4 °C for 25 min. The supernatant was centrifuged again at 13,000x*g* for 15 min, the protein concentration in resulting lysates was measured, and 20 μg of protein from each sample was separated by SDS-PAGE on 4–20% precast polyacrylamide gel (Bio-Rad # 4561096). Western blot analyses were performed according to standard protocols using the following anti-CACNA1A (1:1000, rabbit polyclonal, Abcam) and anti-GAPDH (1:1000, mouse monoclonal, Abcam) antibodies. Equal protein loading was confirmed by Ponceau S staining and normalized for GAPDH immunoreactive band. Detected bands were quantified using ImageJ 1.50i software (National Institutes of Health, Bethesda, MD, USA).

#### Immunofluorescence and lectin fluorescence microscopy

Mouse brains were collected from animals, perfused with 4% PFA in PBS, and postfixed in 4% PFA in PBS overnight. Brains were cryopreserved in 30% sucrose for 2 days at 4 °C, embedded in Tissue-Tek OCT Compound, cut in 40-μm-thick sections, and stored in cryopreservation buffer (0.05 M sodium phosphate buffer, pH 7.4, 15% sucrose, and 40% ethylene glycol) at −20 °C pending immunohistochemistry. The brain sections were washed three times with PBS and permeabilized/blocked by incubating in 5% BSA and 0.3% Triton X-100 in PBS for 1 h at room temperature. Incubation with primary antibodies, rabbit polyclonal anti-aggrecan (1:1000; Millipore, AB1031, RRID:AB_90460) , mouse monoclonal anti-PV (1:2000; Swant, 235, RRID:AB_10000343), rabbit monoclonal anti-mouse GFAP (1:300, DSHB 8-1E7-s), rat monoclonal anti-mouse lysosomal-associated membrane protein 2 (1:200, DSHB ABL-93-s), rabbit polyclonal anti-mouse NeuN (1:250, Millipore Sigma MABN140), rabbit polyclonal anti-mouse CD68 (1:200, Abcam ab125212) and biotinylated Wisteria Floribunda Agglutinin (WFA) to label PNNs (VectorLabs) diluted in 1% BSA and 0.3% Triton X-100 in PBS, was performed overnight at 4 °C. The mouse brain sections were washed three times with PBS and counterstained with Alexa Fluor–labeled secondary antibodies (dilution 1:400) for 2 h at room temperature. After washing three times with PBS, the mouse brain sections were treated with TrueBlack Lipofuscin Autofluorescence Quencher (23007, dilution 1:10; Biotium) for 1 min, and then again washed three times with PBS. The slides were mounted with Prolong Gold Antifade mounting reagent with DAPI (P36935; Invitrogen) and analyzed using a Leica DM 5500 Q upright confocal microscope (10×, 40×, and 63× oil objective, NA 1.4). Images were processed and quantified using ImageJ 1.50i software (National Institutes of Health) in a blinded fashion. Panels were assembled with Adobe Photoshop.
